# Disruption of the interferon-gamma axis limits chimeric antigen receptor T cell efficacy against acute myeloid leukemia

**DOI:** 10.64898/2026.08.28.747900

**Authors:** Nat Murren, Isaac King, Lauren Mahoney, Jarron Roy, Otto A. Kletzien, Madeline Collins, Sarah Geffe, Luke Kalcheim, Rebecca M. Richards

## Abstract

Despite the success of chimeric antigen receptor (CAR) T cell therapy for treatment of B cell acute lymphoblastic leukemia (B-ALL), its translation to acute myeloid leukemia (AML) has been hindered by limited efficacy and significant toxicity. Interferon-gamma (IFNγ) blockade with emapalumab has recently emerged as a promising strategy to mitigate CAR T cell-related toxicities in B cell malignancies, based on evidence that IFNγ is largely dispensable for optimal CAR T cell activity in B-ALL. Whether IFNγ signaling is similarly non-essential in the AML context remains unclear. Here, we demonstrate that disruption of the IFNγ axis impedes anti-AML CAR T cell function and prevents upregulation of target antigen CD123, the apoptotic mediator Fas, and the adhesion molecule ICAM-1 on AML cells. Conversely, exogenous IFNγ enhances CAR T cell cytotoxicity and increases CAR T cell avidity for AML targets. These findings identify IFNγ as a critical mediator of CAR T cell efficacy against AML by promoting increased target antigen expression, enhanced cytotoxicity, and stable CAR T/tumor interactions. Our results suggest that therapeutic IFNγ blockade, including with emapalumab, may compromise CAR T cell responses in AML and should be approached with caution in this disease context.

**Data Sharing Statement:** Bulk RNA-sequencing data analyzed in this study are available in the NCBI Gene Expression Omnibus (GEO) database under accession number GSE159991. All other data generated during this study are available from the corresponding author upon reasonable request.

**Key Points:**

1. IFNγ signaling promotes CAR T cell activity against AML by enhancing antigen expression and immune synapse formation
2. IFNγ blockade may compromise CAR T cell efficacy in AML

## Introduction

CAR T cell therapy has demonstrated remarkable clinical success in treatment of relapsed and refractory B cell malignancies, with high complete response rates and for some patients, durable remissions^1^. Extending CAR T cell therapy to AML has been considerably more challenging, with early phase trials reporting toxicities without meaningful anti-tumor responses^2,3^. As clinical use expands, CAR T cell-related toxicities like cytokine release syndrome (CRS) have proven to be common^4–6^, and can be both life-threatening and dose-limiting. CRS and other related toxicities are typically managed with immunomodulation, often with medications that block specific cytokine axes. Even as cytokine modulation has improved clinical management of CRS^7–10^, systematic identification of strategies that mitigate toxicities without impacting efficacy are needed.

Our current understanding of the cytokine milieu in CAR T cell therapy for leukemia is predominantly derived from pre-clinical studies and clinical observations of CD19 CAR T cell therapy for B cell malignancies^11^. Myeloid and lymphoid blasts both primarily reside in hematopoietic niches that do not pose the same physical microenvironmental barriers of solid tumors, and therefore should be relatively accessible to CAR T cells. However, lineage-specific responses of AML blasts and their contributions to the molecular microenvironment, either at baseline or after interaction with CAR T cells, may induce a suppressive microenvironment that prevents optimal CAR T cell efficacy^12,13^. For example, in a Phase I clinical trial of autologous CD123 CAR T cells for treatment of relapsed and refractory AML (CART-123), high concentrations of soluble factors in patient serum during peak CRS (particularly GM-CSF, IL-3, and FLT3L) led to AML proliferation and resistance to CAR T cell killing^14^.

IFNγ production serves as a benchmark of CAR T cell activity but is also strongly associated with toxicities like CRS^15,16^. In pre-clinical models of B-ALL, disruption of the IFNγ axis decreases toxicity without compromising anti-tumor efficacy^17,18^. Early clinical reports demonstrate that IFNγ antibody emapalumab can mitigate CRS without inhibiting CAR T cell activity against B-ALL^19,20^. In solid tumor models, however, disruption of IFNγ signaling has been associated with increased tumor resistance to CAR T cell killing, partially attributed to reduced expression levels of adhesion molecules like ICAM-1^21^. In AML, ICAM-1 expression plays a key role in sensitizing leukemic blasts to traditional T cell-mediated killing^22,23^. IFNγ also induces the expression of death receptor CD95 (Fas)^24^, which has been shown to enhance CAR T cell activity^25^. Therefore, we hypothesized that unlike in B-ALL, an intact IFNγ axis is required for CAR T cells to effectively kill AML.

To test our hypothesis, we generated model systems to test the impact of pharmacological blockade or genetic deletion of IFNγ on CAR T cell function against both leukemic subtypes. We found that disruption of the IFNγ axis impedes CAR T cell-mediated cytotoxicity against AML but not B-ALL, independent of CAR construct and target antigen. IFNγ blockade specifically prevents upregulation of target antigen CD123, apoptotic mediator Fas, and adhesion molecule ICAM-1 on AML cells. These findings identify IFNγ as a critical mediator of CAR T cell efficacy in AML by promoting both target antigen expression and stable CAR T/tumor interactions. This work identifies fundamental differences between AML and B-ALL susceptibilities to CAR T cells and highlights the importance of defining a tumor-specific molecular microenvironment.

## Methods

### Generation of CAR T cells

Healthy donor-derived T cells were activated with CD3/CD28 Dynabeads (Gibco; Cat#11131D) in AIM-V (Gibco; Cat#12055083) with 100 U/mL of human recombinant IL-2 (Peprotech; Cat#200-02). Cells were transduced on days 2 and 3 on Retronectin (Takara; Cat#T100B) coated plates and Dynabeads were removed on day 4. On days 9-12 post-activation, cells were used in functional assays or cryopreserved. For IFNγ knockouts, Dynabeads were removed on day 2, and T cells were electroporated with 3 sgRNA’s (Genscript) targeting human IFNγ (AAAGAGTGTGGAGACCATCA, TTTCAGCTCTGCATCGTTTT, CCAGAGCATCCAAAAGAGTG) and SpCas9 nuclease (IDT; Cat#1081058) using the P3 4D-Nucleofector Kit (Lonza; Cat#V4XP-3032). Cells were transduced on days 3 and 4. Knockout efficiency was determined by intracellular flow cytometry and sequencing of PCR amplicons (Plasmidsaurus) was analyzed with CRISPResso2^26^ software.

### Generation of Recombinant Leukemia Cell Lines

Leukemia lines were cultured in complete RPMI (+10% FBS + 1% penicillin/streptomycin + 10 mM HEPES) then transduced with retrovirus containing CD123 or CD19. Cells were sorted based on antigen density using the BD FACS Aria. Single-cell clones were expanded over the course of 2-3 weeks then validated with flow cytometry.

### Flow Cytometry

Cells were washed with FACS buffer (PBS + 2% FBS), incubated with human Fc block (BD Biosciences; Cat# 564220) for 15 min, then stained with antibodies suspended in Brilliant Stain Buffer (BD Biosciences; Cat# 563794). CAR expression was detected with anti-mouse IgG F(ab’) (Jackson ImmunoResearch; Cat#115-606-072). PE-Quantibrite beads (BD Biosciences; Cat# 340495) were used to calculate the number of surface molecules per cell from a standard curve. For intracellular staining, samples were stained for surface markers then fixed, permeabilized, and stained for intracellular markers with a FIX and PERM cell permeabilization kit (Invitrogen; Cat# GAS003). All antibodies are listed in Table S1. Samples were run on the Attune NxT cytometer and analyzed using FlowJo software.

### Cytokine Production

Co-culture supernatant was diluted 1:20 and IFNγ or Granzyme B production was measured using ELISA MAX detection kits (Biolegend; Cat#430116 and #439204). Luminescence was measured using the ClarioStar Plus at 450 and 570 nm.

### Incucyte Cytotoxicity Assays

Leukemia cell lines were co-cultured with mock or CAR T cells at various effector:target (E:T) ratios in complete RPMI and imaged every 4h with an Incucyte live cell imager. IFNγ was pharmacologically blocked with 5 μg/mL of anti-human IFNγ (BD Biosciences; Cat#554698). ICAM-1 was blocked with 10 µg/mL Ultraleaf anti-human CD54 (Biolegend; Cat#322722). Anti-IgG1 isotype was used as a control (BD Biosciences; Cat#554721). For serial stimulation assays, additional tumor cells were added every 24h and antibodies were replenished every 48h. Tumor cell growth based on GFP fluorescent intensity and normalized to initial values on day 1, or with each re-stimulation.

### Lactate Dehydrogenase (LDH) Cytotoxicity Assays

AML and B-ALL patient samples were co-cultured with CD123 (AML) or CD19 (B-ALL) CAR T cells for 48 h. LDH was quantified using an LDHCytox Assay (Biolegend; Cat#426401). High and low controls were mock T cells co-cultured with the corresponding primary sample and lysed with the provided lysis buffer (high) or cultured alone (low). Absorbance of samples was measured at 490 nm using the ClarioStar Plus. Primary leukemia samples are accessible under IRB # 2025-0727.

### Phenotyping Leukemia after Exposure to Conditioned Media

Mock- or CAR-transduced T cells were co-cultured with tumor cells 1:1 for 24h. Supernatants were collected, centrifuged, and filtered to remove any remaining cells. Fresh tumor cells were washed with PBS and then incubated with the conditioned media for 24h. Following incubation, cells were harvested and subjected to flow cytometry.

### Avidity Studies

Tumor cells were cultured in complete RPMI with or without 10 ng/mL recombinant human IFNγ (Peprotech; Cat# 300-02) for 24h at 37°C. Tumor cells were seeded onto a z-Movi chip and CAR T cells were labeled with CellTrace™ Far Red (Thermo Fisher; Cat# C34564) per manufacturer’s protocol. For avidity measurements, CAR T cells were added to tumor cell-coated chips, incubated for 5 min, subjected to increasing acoustic force using the z-Movi, then analyzed using z-Movi Control and Analysis Software.

### Statistical Analysis

Statistical analyses were carried out using GraphPad Prism software or R Studio. P values were calculated with statistical tests described in each figure legend and are denoted with asterisks as follows: P > 0.05 = not significant (ns), P </= 0.05 = *, P < 0.01 = **, P < 0.001 = ***, P < 0.0001 = ****.

## Results

### IFNγ blockade impedes optimal CAR T cell killing of AML but not B-ALL

To determine whether IFNγ is necessary for CAR T cell cytotoxicity against AML, we performed *in vitro* cytotoxicity assays in the presence of isotype control or anti-IFNγ (αIFNγ) antibody. We compared cytotoxicity of CD19 CAR T cells against B-ALL cell line Nalm-6 or CD123 CAR T cells against AML cell lines THP-1 or OCI-AML3. To mimic the repeated CAR T/tumor interactions that occur in patients, we implemented a serial challenge model that included re-stimulation with tumor cells every 24h for 3 days (Fig 1A). To ensure that pharmacological blockade of IFNγ alone did not impair tumor cell growth, we confirmed that neither Nalm-6 nor THP-1 proliferation was inhibited with increasing concentrations of αIFNγ (Fig S1A-B). 5 µg/mL was sufficient to effectively block IFNγ (Fig S1C). To determine the duration of IFNγ blockade for longer-term *in vitro* assays, we measured IFNγ concentration every 24h for 4 days. IFNγ blockade was sustained for the first 48h, but the effect was lost by 72h. Notably, αIFNγ had no impact on the secretion of Granzyme B (Fig S1D). Based on these results, αIFNγ was added at initial co-culture and supplemented every 48h for serial stimulation assays (Fig 1A).

**Figure 1.**
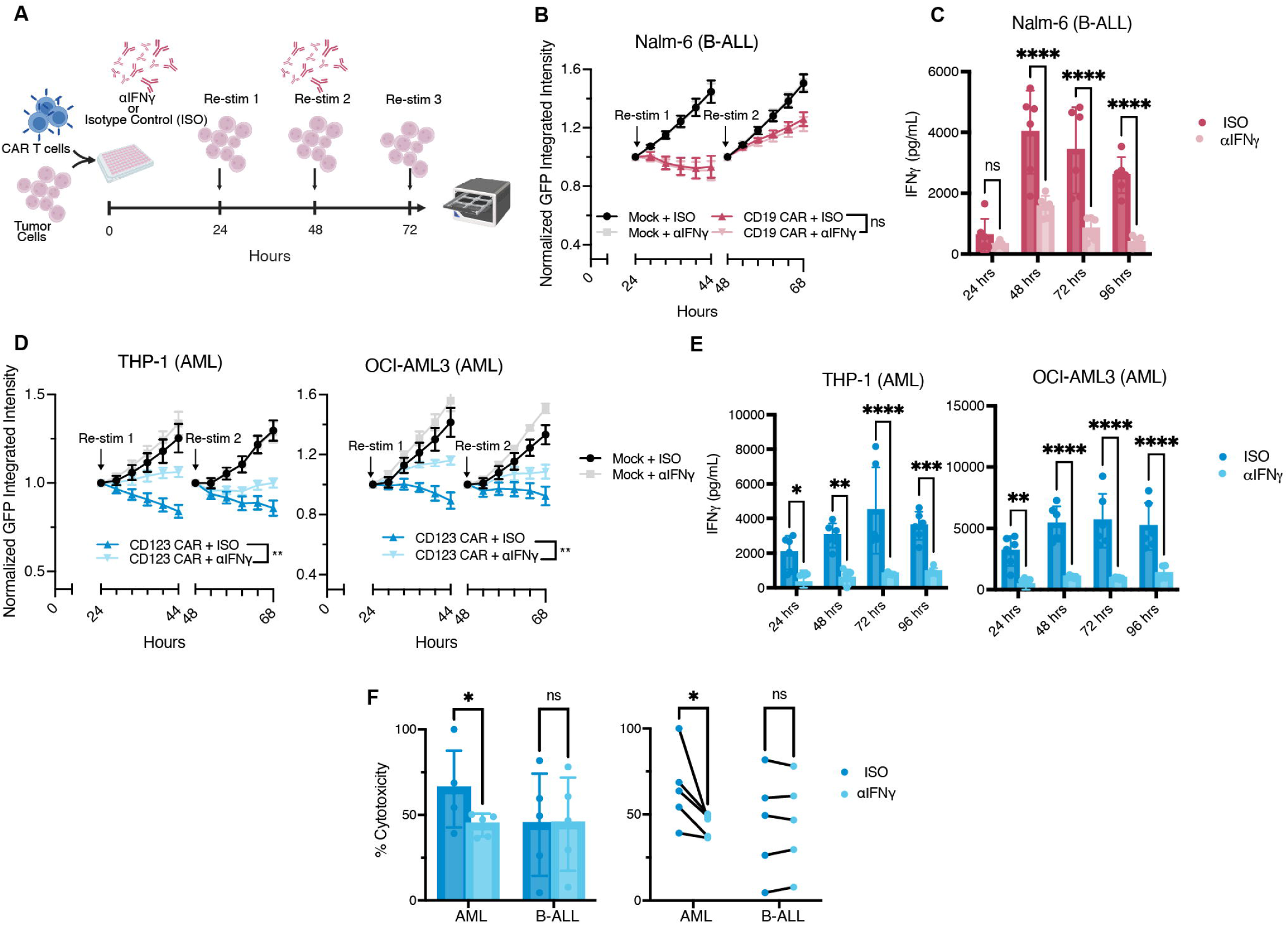
IFNγ blockade impedes optimal CAR T cell cytotoxicity against AML. **A.** Schematic of the serial stimulation cytotoxicity assay workflow. **B.** Co-culture of Nalm-6 B-ALL cells with CD19 CAR T cells in the presence of isotype control antibody or anti-IFNγ antibody (αIFNγ). Tumor growth was quantified as relative GFP intensity using Incucyte® imaging and re-normalized every 24h following each tumor re-challenge. Data from n=3 T cell donors are shown for two 24h periods (after re-stimulation #1 and #2) at an E:T ratio of 1:4. Statistics shown are based on differences between CAR+ISO and CAR+αIFNγ during the 48 hour period shown (Welch’s t-test of AUC). Full time courses and individual donors are shown in Fig S2. **C.** IFNγ concentrations in Nalm-6/CD19 CAR co-culture supernatants measured by ELISA every 24h over 96h at a 1:4 E:T ratio, n=3 T cell donors (two-way ANOVA). **D.** Co-culture of AML cell lines THP-1 (right) and OCI-AML3 (left) with CD123 CAR T cells with isotype control or αIFNγ. Tumor growth and statistics were assessed as in B, also at a 1:4 E:T ratio, n=3. Full time courses and individual donors are shown in Fig S3. **E.** IFNγ concentration measured by ELISA from supernatant from CD123 CAR T/THP-1 or CD123 CAR T/OCI-AML3 cocultures over 96h at a 1:4 E:T ratio, n=3 T cell donors (two-way ANOVA). **F.** LDH-based cytotoxicity assay of primary patient-derived AML (n=5) or B-ALL (n=5). Bar plots show mean percent cytotoxicity relative to mock T cell co-culture controls in the presence of isotype control or αIFNγ (left). Paired plots connect individual primary leukemia samples across treatment conditions (right). Statistics based on change in percent cytotoxicity between ISO and αIFNγ for matched leukemia donors (paired t-test).

Consistent with previous studies^17,18^, CD19 CAR T cells retained their ability to kill B-ALL Nalm-6 cells with or without αIFNγ, even after three consecutive re-challenges when anti-leukemic effect began to wane (Fig 1B, S2A). We demonstrated significant, though not complete, reduction of IFNγ throughout the serial stimulation assay (Fig 1C). However, IFNγ blockade significantly reduced tumor cell killing in CD123 CAR T/AML co-cultures over time compared to the isotype control (Fig 1D, S2A-B). We confirmed that IFNγ blockade did not affect Granzyme B secretion throughout the serial stimulation assay (Fig 1E, S3C).

To validate the differential IFNγ-dependence of CAR T cell activity against clinically representative leukemia samples, we co-cultured CD123 or CD19 CAR T cells with primary AML or B-ALL patient samples, respectively, with αIFNγ or isotype control antibody. All AML samples expressed CD123 and all B-ALL samples expressed at CD19, albeit at varying intensities (Fig S4A-B). An LDH assay quantified cytotoxicity of CAR T cells relative to donor-matched mock T cells, at E:T ratios chosen to avoid signal oversaturation (1:1 for CD123 CAR T/AML and 1:8 for CD19 CAR T/B-ALL). The relative CAR T cell cytotoxicity was significantly reduced in primary AML cultures treated with IFNγ blockade compared to control antibody, but this difference was not seen in primary B-ALL co-cultures, aligning with immortalized cell lines (Fig 1F). Together, these data confirm that blockade of IFNγ impairs CAR T cell efficacy against AML but not B-ALL.

### CAR T cell reliance on IFNγ in AML is independent of scFv-target antigen pairing

We next wanted to compare the differential impact of IFNγ blockade in a CAR- and target antigen-independent manner. We engineered Nalm-6 B-ALL cells to constitutively express AML target antigen CD123 (N6-123), and AML cell lines to express B-ALL target antigen CD19 (Fig 2A). Using single-cell cloning, we identified N6-123 cell lines with low- and medium-expression that most closely matched endogenous AML antigen density and were therefore used for downstream functional assays (Fig 2B, Fig S5A). We then evaluated CD123 CAR T cell cytotoxicity against AML or N6-123 cell lines in serial stimulation assays, with and without IFNγ blockade. When co-cultured with N6-123 cells, CD123 CAR T cells displayed no significant differences in tumor cell killing in the presence of αIFNγ compared to the control (Fig 2C, Fig S5B). Confirming effective blockade, IFNγ was significantly reduced in αIFNγ-treated co-cultures compared to controls, without significant differences in Granzyme B secretion (Fig. S5C-D). We extended this approach to AML models by engineering THP-1 and OCI-AML3 cell lines to express CD19 at a similar antigen density to native CD19 expression on Nalm-6 cells (Fig 2D, S6A). Blocking IFNγ limited CD19 CAR T cell cytotoxicity toward CD19-expressing AML cells (Fig. 2E, S6B-D). Therefore, IFNγ dependence appears to be driven by tumor-intrinsic properties of AML rather than by the specific CAR or target antigen.

**Figure 2.**
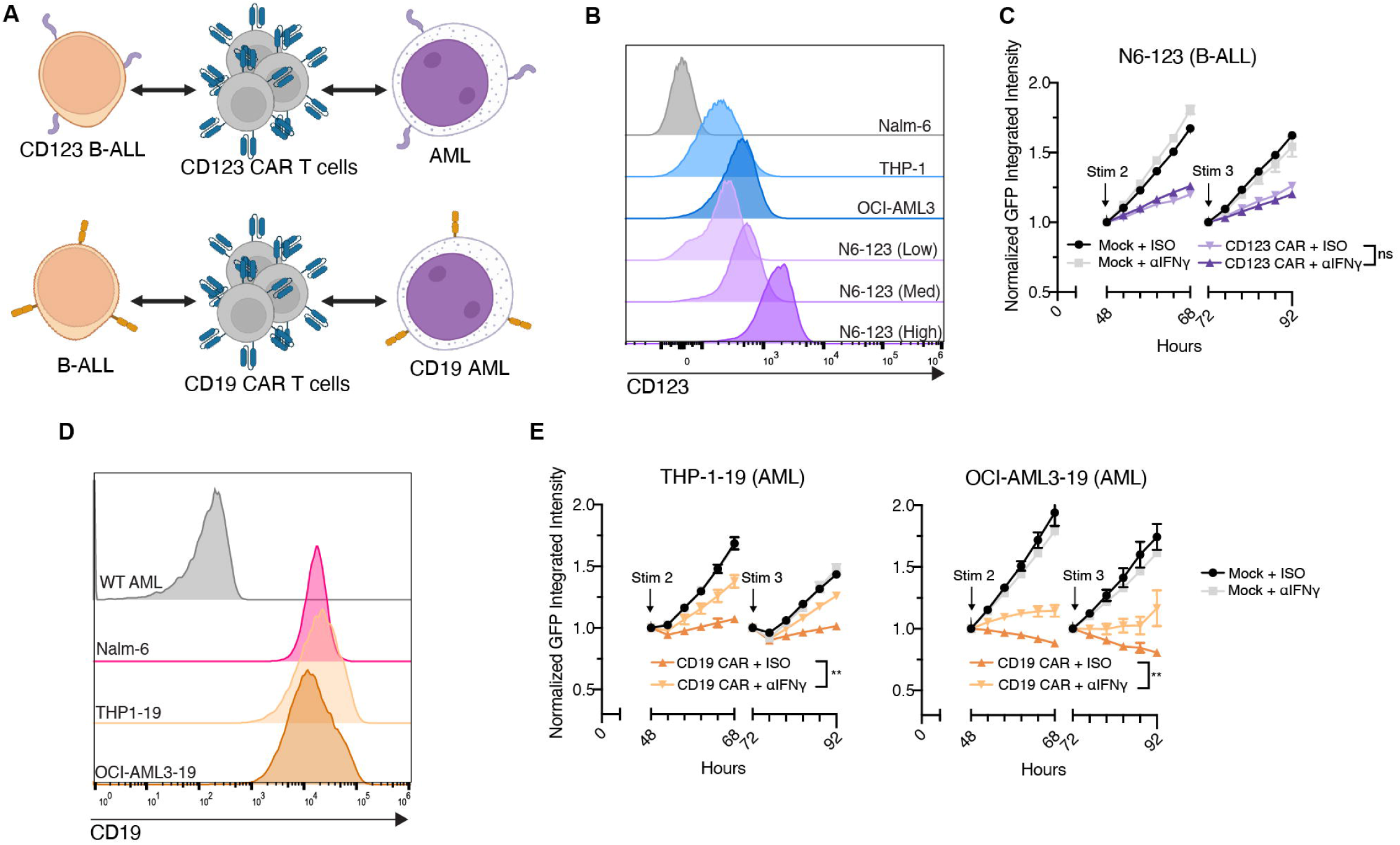
IFNγ blockade reduces CAR T cell mediated killing of AML independent of target antigen. **A.** Schematic illustrating engineered CD123-expressing Nalm-6 (N6-123) and CD19-expressing AML cell lines (THP1-19 and OCI-AML3-19). **B.** Flow cytometry analysis of CD123 expression on AML cell lines (THP-1 and OCI-AML3) and single cell selected N6-123 cell line. **C.** Serial stimulation co-culture of N6-123 cells and CD123 CAR T cells with isotype control or with αIFNγ. Tumor growth and statistics were assessed as in Figure 1, with data from n=3 T cell donors shown for two 24h periods (after re-stimulation #2 and #3) at a 1:4 E:T ratio. Full time courses and individual donors are shown in Fig S5. **D.** Flow cytometry analysis of CD19 expression on Nalm-6 and single cell selected CD19 AML cell lines. **E.** Serial stimulation co-culture of THP1-19 (left) and OCI-AML3-19 (right) cells and CD19 CAR T cells with isotype control or αIFNγ. Tumor growth and statistics were assessed as in Figure 1, with representative donor data at a 1:4 E:T ratio. Full time course and individual donors shown in Fig S6.

### Genetic knockout of IFNγ impedes CAR T cell killing of AML but not B-ALL

While IFNγ blockade mimics the mechanism of action of emapalumab, pharmacological blockade did not completely abrogate IFNγ. Because AML cells can be induced to secrete IFNγ in some settings, we wanted to isolate the IFNγ depletion to CAR T cell production and secretion. We used CRISPR-Cas9 to generate IFNγ knock out (IFNγKO) CD123 CAR T cells (Fig. S7A). Effective knockout was confirmed with intracellular flow cytometry (Fig. S7C) and PCR amplicon sequencing^26^ (Fig S7D). Wild type (WT) and IFNγKO CD123 CAR T cells had comparable expansion after electroporation (Fig S7B), exhibited similar CD4/8 ratios (Fig. S7E), and displayed similar surface expression of markers associated with CAR T cell exhaustion and dysfunction, including LAG3, TIM3, and PD-1 (Fig. S7F).

WT or IFNγKO CD123 CAR T cells were then subjected to our serial stimulation assay at low E:T ratios (Fig 3A). IFNγKO CAR T cells had near-complete and sustained abrogation of IFNγ secretion compared to WT CAR T cells for up to 96h, without affecting Granzyme B production (Fig 3C-D). Consistent with IFNγ blockade, IFNγKO CD123 CAR T cells exhibited reduced cytotoxicity against AML compared to WT CD123 CAR T cells (Fig 3B, S8A). In contrast, there was no difference in WT and IFNγKO CD123 CAR T cell killing of engineered N6-123 cells, validating that IFNγ is dispensable for optimal CAR T cell efficacy against B-ALL but critical in the AML context (Fig 3E-G, S8B).

**Figure 3.**
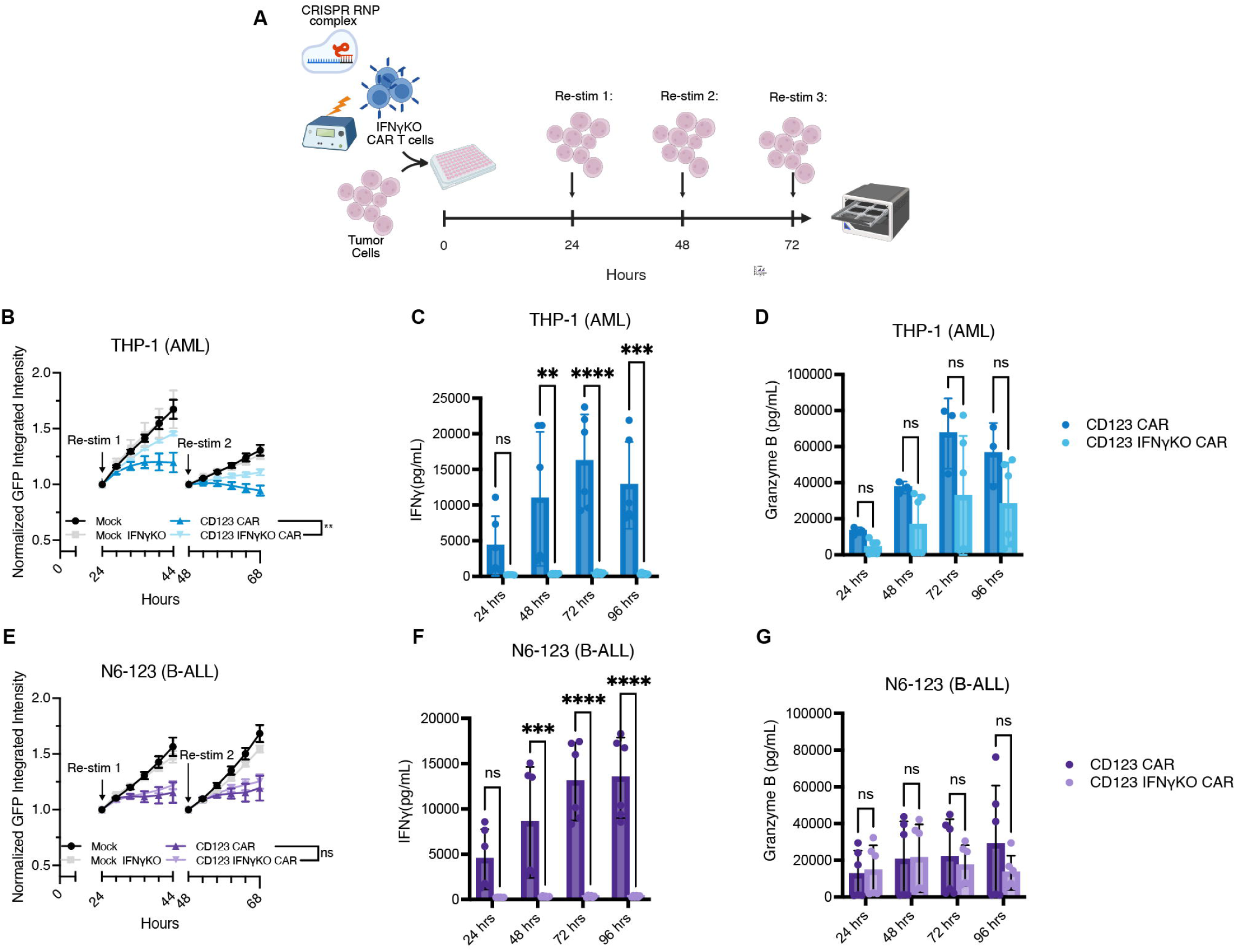
Genetic knockout of IFNγ reduces CAR T cell mediated killing of AML but not B-ALL. **A.** Schematic of the CRISPR KO serial stimulation cytotoxicity assay workflow. **B.** Serial stimulation co-culture of N6-123 cells and wild-type (WT) or IFNγ knockout (IFNγKO) CD123 CAR T cells. Tumor growth was assessed as in Figure 1 and 2, with summary donor data n=3 at a 1:2 E:T ratio (Welch’s t-test of AUC). Full time course and individual donors shown in Fig S8. **C-D.** ELISA quantification of IFNγ (C) and Granzyme B (D) in supernatant from N6-123 and CD123 WT or KO CAR cocultures every 24h for 96 h, n=3 (two-way ANOVA) **E.** Serial stimulation co-culture of CD19 THP-1 CD123 CAR T cells with or without IFNγKO. Full time course and individual donors shown in Fig S8. **F-G.** ELISA quantification of IFNγ (C) and Granzyme B (D) in supernatant from THP-1 and CD123 WT or KO CAR cocultures every 24h for 96 h, n=3 (two-way ANOVA).

### IFNγ blockade limits upregulation of CD123 on AML

CD123 cell surface expression increases in response to pro-inflammatory cytokines TNFα and IFN ^27^. We hypothesized that this dynamic expression was a primary driver of IFNγ dependence in AML, and that interrupting the IFNγ axis prevents CD123 upregulation. We measured CD123 expression levels on AML during our serial stimulation assays, with and without IFNγ blockade. Indeed, THP-1 AML target cells upregulated CD123 after 48h or 96h of co-culture with CD123 CAR T cells, when compared to co-culture with mock T cells. Of note, this experiment was designed with a low E:T ratio of 1:16 to prevent rapid cytotoxicity of all target cells. Confirming our hypothesis, addition of αIFNγ prevented CD123 upregulation (Fig 4A). In the presence of IFNγ blockade, CD123 upregulation was minimal (Fig. 4B). Exposure to supernatant from CAR T cell/AML co-cultures increased CD123 expression in comparison to the mock T cell control, and the addition of IFNγ blockade diminished this effect (Fig 4C), confirming that this phenomenon is driven primarily by secreted IFNγ rather than by cell-to-cell interactions. In contrast to AML, CD123 expression was stable on N6-123 B-ALL cells across all conditions and there was no difference in quantitative antigen density with or without IFNγ blockade (Fig S9A-C).

We also confirmed a lack of dynamic target antigen expression for CD19. THP1-19 and Nalm-6 cells collected from co-cultures had similar CD19 expression at baseline, with isotype control, or with αIFNγ (Fig. S10A-B). Therefore, while CD123 dynamic expression may contribute to the differential dependence on IFNγ observed between AML and B-ALL models, it is unlikely the sole mechanism underlying this phenomenon, as CD19 expression was not similarly modulated.

**Figure 4.**
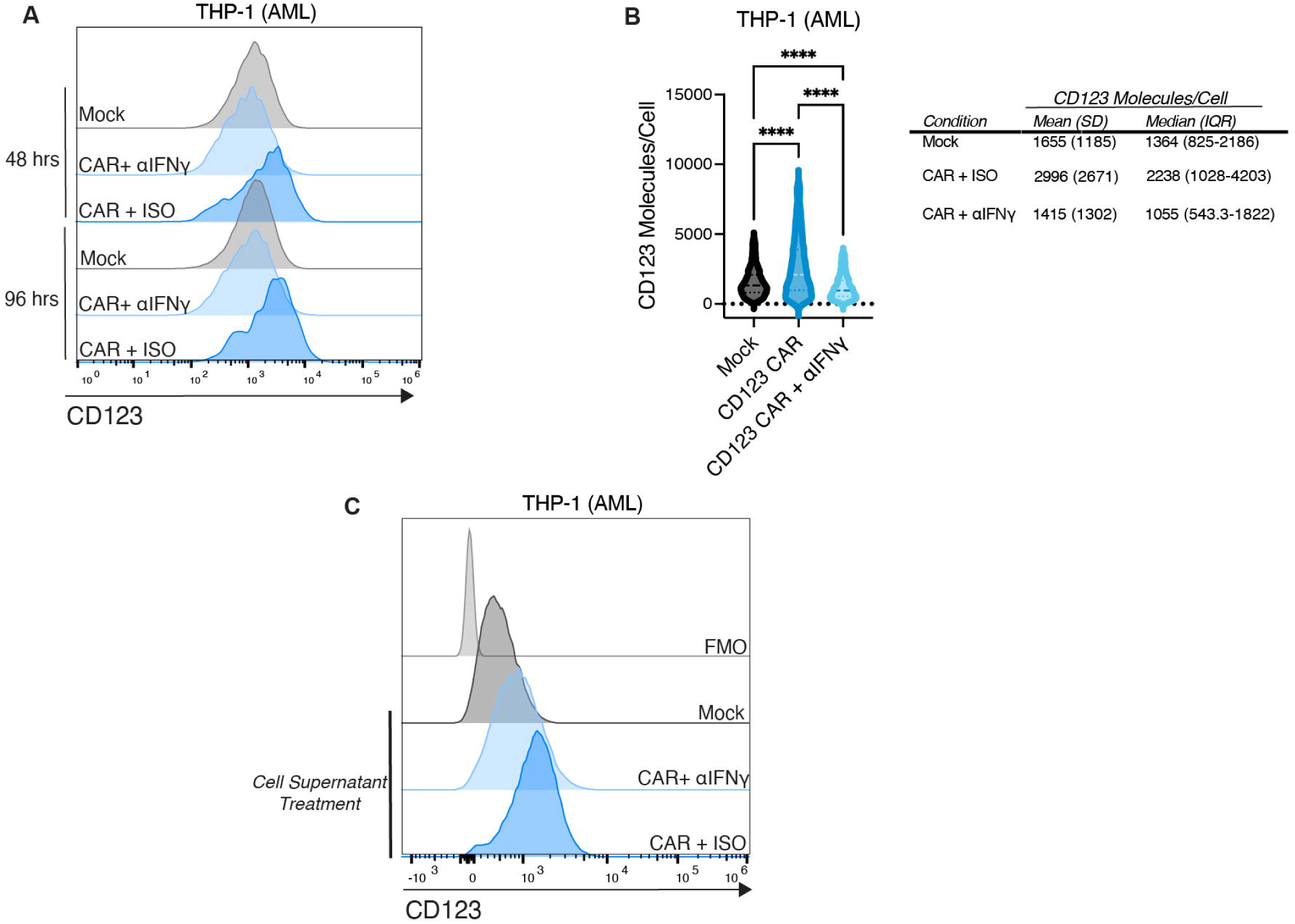
Upregulation of CD123 in the context of CAR T cell inflammation correlates with improved CAR T cell killing of AML. A-B. Flow cytometry of CD123 expression on THP-1 cells at 48 and 72h during a serial stimulation assay, with or without αIFNγ treatment at a 1:16 E:T ratio represented as (A) relative intensity by flow cytometry, (B) molecules per cell calculated from a standard curve generated with BD Quantibrite beads, measured after 48h of co-culture with mock or CAR T cells. Data from 1×10^4^ individual cells shown with statistics based on mean difference between CAR+ISO and CAR+αIFNγ (one-way ANOVA). **C.** CD123 expression on THP-1 cells exposed to supernatant from THP-1/Mock, THP-1/CAR, or THP-1/CAR+αIFNγ co-cultures for 24 h.

### IFNγ blockade mitigates upregulation of key surface molecules on AML

We next investigated whether IFNγ blockade influenced other cell surface molecules that are known to be important for CAR T cell activity, and whether there are differences in AML and B-ALL. We reanalyzed a previously described RNA-seq dataset^28^ and identified several genes that were differentially expressed across three AML cell lines after IFNγ exposure compared to baseline expression. Death receptor CD95 (Fas) and adhesion molecule ICAM-1 (Fig S11A), both previously identified as important enhancers of CAR T cell cytotoxicity^21,25,29^, were upregulated in IFNγ-primed AML. We therefore hypothesized that IFNγ blockade in the AML setting could prevent the ICAM-1 and Fas upregulation necessary for optimal CAR T cell cytotoxicity.

To assess the effect of CAR T cell related inflammation and of IFNγ blockade on Fas and ICAM-1 expression on tumor cells, we measured cell surface expression after addition of conditioned media from AML/CD123 CAR T or N6-123/CD123 CAR T co-cultures. Fas was upregulated on AML cells exposed to CAR T/AML but not mock T/AML conditioned media. Addition of αIFNγ limited upregulation of both ICAM-1 and Fas on AML cells. In contrast, Fas expression remained unchanged on N6-123 cells after exposure to N6-123/CAR T cell supernatant, irrespective of IFNγ blockade (Fig. 5A). Notably, this pattern was preserved in cross-conditioning experiments, where N6-123 cells maintained consistent Fas expression after exposure to CAR T/AML conditioned media and AML cells upregulated Fas when exposed to CAR T/B-ALL cell conditioned media, reflecting an AML-intrinsic responsiveness to IFNγ blockade (Fig. S11B).

**Figure 5.**
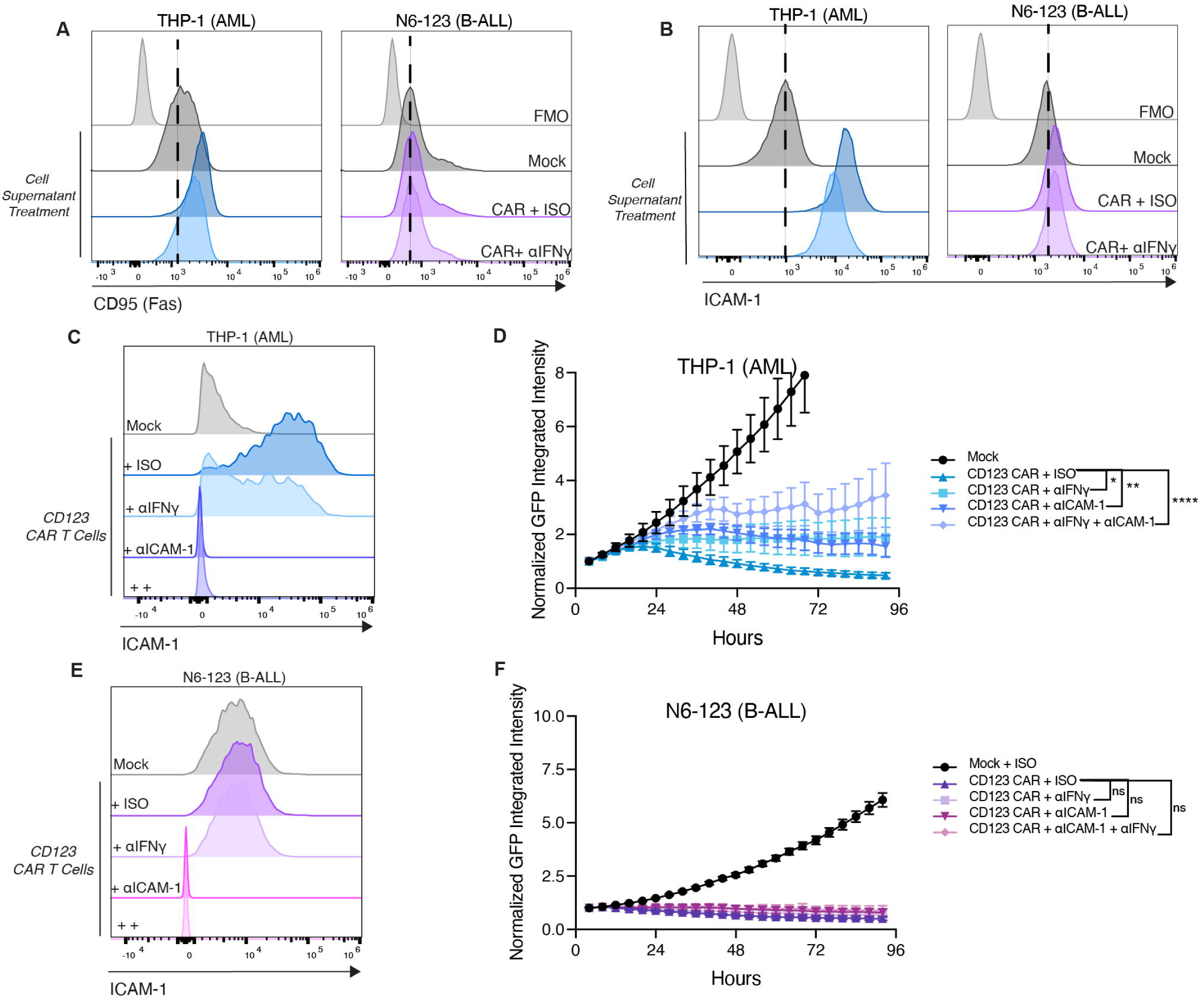
IFNγ blockade modulates Fas and ICAM-1 on AML. **A-B.** Flow cytometry of (A) CD95 (Fas) and (B) ICAM-1 expression on AML and B-ALL tumor cell lines after 24h of exposure to cell line-matched conditioned media. Dotted lines are drawn based on peak fluorescence levels of controls exposed to mock/tumor conditioned media. **C.** Flow cytometry of ICAM-1 expression on THP-1 AML cells after 24-hour co-culture with CD123 CAR T cells with addition of isotype control, αIFNγ, anti-ICAM-1 (αICAM-1) or αIFNγ and αICAM-1 in combination at a 1:8 E:T ratio. **D**. Incucyte cytotoxicity assay of CD123 CAR T cells co-cultured with THP-1 cells for 96h with the antibody conditions described above at a 1:2 E:T ratio. Tumor growth was quantified as relative GFP intensity normalized to values at 4 h. Shown is summary data with statistics based on comparisons between each CAR T cell condition, n=3 T cell donors (one-way ANOVA of AUC). **E.** Flow cytometry of ICAM-1 expression on N6-123 B-ALL cells after 24-hour co-culture with CD123 CAR T cells with addition of isotype control, αIFNγ, αICAM-1, or αIFNγ and αICAM-1 in combination at a 1:8 E:T ratio. **F**. Incucyte cytotoxicity assay of CD123 CAR T cells co-cultured with N6-123 cells for 96h with the same antibody conditions described above at a 1:2 E:T ratio. Tumor growth and statistics quantified as in Fig 5D, n=3 T cell donors (one-way ANOVA of AUC).

AML cells also displayed increased ICAM-1 surface expression following exposure to both CAR T/AML and CAR T/B-ALL conditioned media. However, this expression was only partially reduced by the addition of IFNγ blockade, suggesting that ICAM-1 regulation on AML relies on a combination of inflammatory factors. In contrast, ICAM-1 expression on N6-123 cells, though slightly upregulated when exposed to CAR T/tumor conditioned media, was not altered by the presence or absence of αIFNγ (Fig 5B, S11C).

To determine whether blockade of the IFNγ-ICAM-1 axis would be sufficient to impact CAR T cell cytotoxicity, we assessed CD123 CAR T cell killing of AML or B-ALL in the presence of IFNγ and/or ICAM-1 blockade. Following a 24h co-culture with CD123 CAR T cells, ICAM-1 expression on THP-1 cells increased relative to co-culture with mock T cells, and IFNγ blockade diminished this effect. However, expression levels were still higher than mock control, αICAM-1 alone, and combined blockade (αIFNγ + αICAM-1, Fig 5C). In cytotoxicity assays, ICAM-1 or IFNγ blockade independently decreased CAR T cell killing of AML, but the combination resulted in the most pronounced impairment of cytotoxicity (Fig 5D). In contrast, ICAM-1 expression on N6-123 cells after co-culture with CD123 CAR T cells remained unchanged across all experimental conditions (Fig 5E), and CD123 CAR T cell cytotoxicity against N6-123 cells was not significantly affected by ICAM-1 blockade, IFNγ blockade, or their combination (Fig 5F).

### Exogenous IFNγ enhances CAR T cell cytotoxicity and avidity against AML

Since disruption of IFNγ reduced CAR T cell killing of AML, we wanted to see whether exogenous IFNγ could boost CAR T cell activity against AML. We supplemented CD123 CAR T/AML or CD123 CAR T/B-ALL co-cultures with exogenous IFNγ (10 ng/mL), then assessed cytotoxicity over 72h. CD123 CAR T cell cytotoxicity against AML increased in the presence of supplemental IFNγ (Fig 6A). However, exogenous IFNγ did not improve cytotoxicity against N6-123 cells even at low E:T ratios of 1:32 at which the CAR T cells begin to fail to control tumor (Fig 6B). Additionally, improved cytotoxicity toward AML correlated with an expected increase in the expression of key surface molecules, including CD123, Fas, and ICAM-1 (Fig 6C). Again, IFNγ-dependent dynamic expression of these molecules was not seen on N6-123 cells. Finally, IFNγR1 surface expression was downregulated on all cell lines following 24h of IFNγ exposure, consistent with active IFNγ signaling across both leukemic subtypes (Fig. S12).

**Figure 6.**
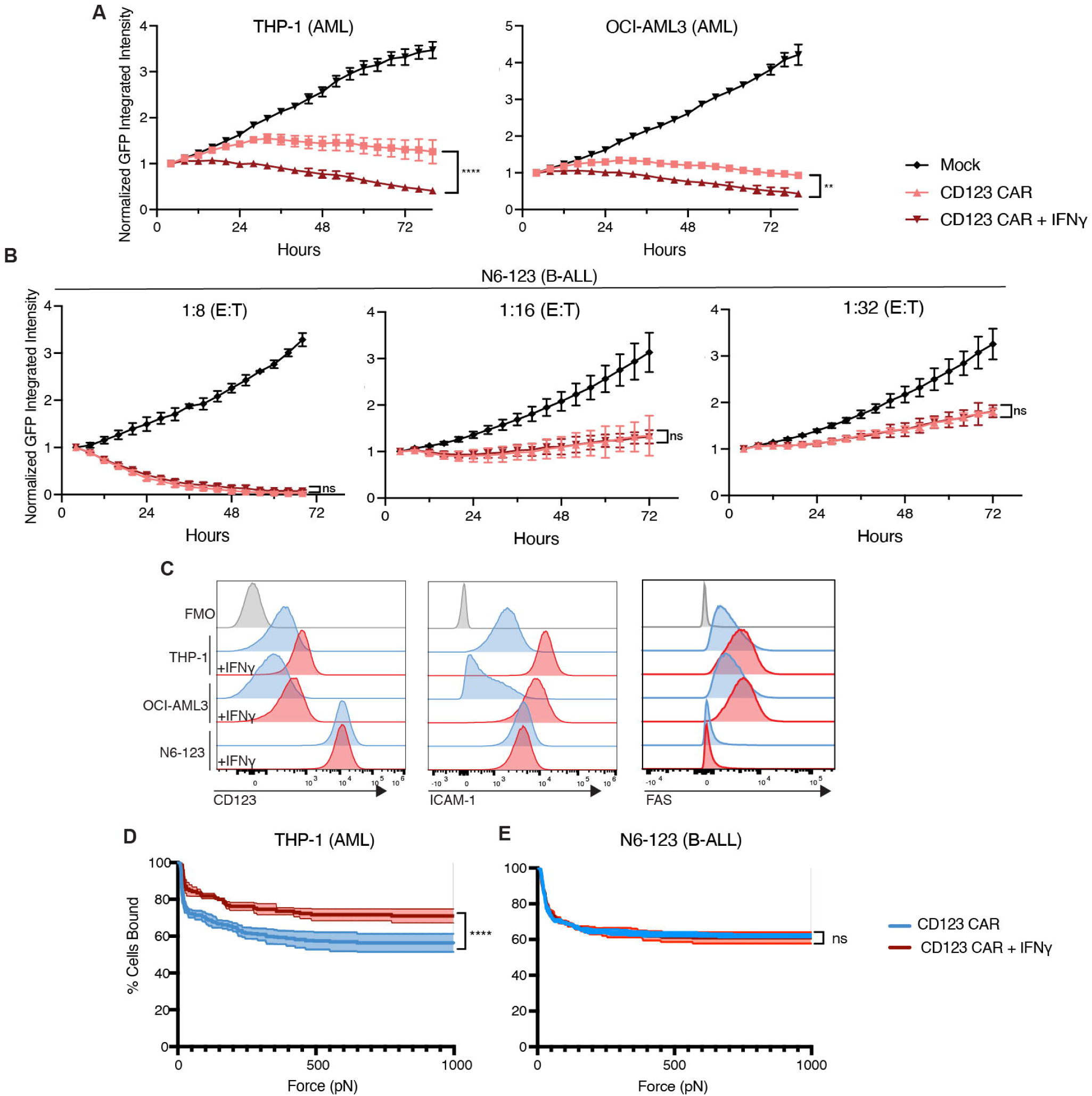
Exogenous IFNγ upregulates ICAM-1 and Fas and enhances anti-AML CAR T cell cytotoxicity and avidity. **A-B**. Incucyte cytotoxicity assays of CD123 CAR T cells co-cultured with (A)THP-1 and OCI-AML3 cells at a 1:8 E:T ratio or (B) N6-123 cells at a range of 1:8-1:32 E:T ratios, alone or in combination with 10 ng/mL of IFNγ for 72 h. Tumor growth was quantified as relative GFP intensity normalized to values at 4 h. Statistics based on comparisons between each CAR T cell condition, n=3 T cell donors (Welch’s t-test of AUC). **C.** Flow cytometry of CD123 (left), Fas (middle), and ICAM-1 (right) expression on AML and B-ALL cell lines after 24-hour pre-incubation with 10 ng/mL of IFNγ **D**-**E.** Z-movi avidity analysis of CD123 CAR T cells to (D) THP-1 and (E) N6-123 cells alone and pre-conditioned in 10 ng/mL of IFNγ for 24 h. Plots show average percentage of T cells bound (y-axis) over increasing acoustic force applied (x-axis) ranging from 0-1000 pN after initial 5-minute incubation on a target cell monolayer. Statistical comparisons performed using Welch’s t-test of AUC.

Since exogenous IFNγ enhanced both target antigen CD123 and adhesion molecule ICAM-1 expression on AML, we next investigated whether this similarly correlated with an increase in CAR T cell avidity. THP-1 and N6-123 cells were pre-incubated with IFNγ for 24h to induce expression of CD123, ICAM-1, and Fas, then exposed to CD123 CAR T cells in an acoustic force-based avidity assay. A greater percentage of CD123 CAR T cells remained bound to IFNγ-treated THP-1 cells compared to untreated THP-1 cells, indicating IFNγ-dependent avidity enhancement (Fig 6D). In contrast, no significant difference in CAR T cell avidity was observed for N6-123 cells with or without IFNγ pre-incubation (Fig 6E).

## Discussion

For clinical CAR T cell indications to expand beyond B cell malignancies, and to design rational strategies for toxicity management that do not interfere with efficacy, it is critical to dissect the molecular and cellular microenvironment with a tumor-specific perspective. Here, we generated a reductionist model system to directly compare IFNγ dependency in AML and B-ALL to test our hypothesis that IFNγ-specific influences were distinct in these two hematologic malignancies.

Broadly, IFNγ signaling exerts context-dependent effects on AML biology, influencing both proliferative capacity and immune sensitivity depending on the therapeutic pressure applied. Intra-leukemic IFNγ signaling is associated with cell cycle suppression and reduced expansion of AML patient samples contributing to chemoresistance^30,31^. However, in the context of T cell-mediated immunotherapies, elevated IFNγ signaling in AML was a strong predictor of therapeutic response to bispecific T cell engagers, mediated in part by MHC-II upregulation and STING pathway activation^32–34^. Our study extends this concept to CAR T cell therapy.

We confirmed that disrupting the IFNγ axis does not affect CAR T cell activity against B-ALL, as has been previously shown^17,18^. In contrast, pharmacologic or genetic disruption of the IFNγ axis impaired anti-AML activity, suggesting that IFNγ signaling is critical for optimal CAR T cell function in AML. This is of particular interest for CD123 CAR T cells, as CD123 is a promising AML target antigen, but is expressed at or near the antigen density threshold for CAR T cell recognition^35^, and is dynamically regulated in a pro-inflammatory microenvironment^36^. However, AML-specific IFNγ-dependence also extended to the constitutively and stably expressed target antigen CD19, suggesting a role for other mechanisms.

To determine the importance of other key IFNγ-driven regulatory pathways, we also assessed Fas and ICAM-1 expression in our model systems as these are two cell surface molecules expressed on a variety of tumors and known to be important to CAR T cell function in some contexts^21,25,29,37^. The Fas/FasL death receptor pathway is an alternative killing pathway to perferon/granzyme-mediated cytotoxicity, where FasL expressing T cells induce apoptosis in Fas expressing target cells upon interaction^38^. In CAR T cell therapy, Fas expression on tumor cells can facilitate apoptosis of nearby target antigen negative “bystander” cells, which would be desirable to enhance anti-tumor effect for heterogeneous malignancies like AML^39,40^. ICAM-1 is an essential component of the traditional TCR immune synapse formation^41^, and genetic knockout of ICAM-1 in AML reduces its sensitivity to native T cell killing^23^. Fas and ICAM-1 expression were regulated on AML, but not B-ALL, in an IFNγ-dependent manner, and increased expression correlated with better CAR T cell killing. Even independent of IFNγ blockade, ICAM-1 blockade compromised anti-AML but not anti-B-ALL CAR T cell cytotoxicity, indicating an AML-specific dependence that mirrors solid tumor models^21^. Therefore, we propose that CAR T cell engineering and microenvironmental remodeling strategies in AML should be distinct from that of B-ALL. Additionally, this work highlights Fas and ICAM-1 as independent targets to improve CAR T cell efficacy in AML.

Our findings have important implications for the clinical use of cytokine-modulating therapies in AML CAR T cell therapy. Emapalumab has shown promise for mitigating CRS, particularly in B-cell malignancies. However, our results suggest that IFNγ plays a pivotal role in promoting CAR T cell activity against AML. As a result, IFNγ blockade in this setting could unintentionally reduce CAR T cell efficacy by limiting IFNγ-dependent sensitization of AML cells to immune killing.

Additionally, therapeutic strategies augmenting IFNγ signaling may further enhance CAR T cell efficacy against AML. CAR T cells that can locally modulate cytokine signaling have previously been proposed as a method of overcoming the immunosuppressive microenvironment of solid tumors^42,43^ and may be a viable approach to transiently increase IFNγ in hopes of improving therapeutic response against AML. Of course, any attempt to therapeutically exploit this pathway will require careful balancing of anti-leukemic benefit with the potential for supratherapeutic systemic IFNγ that could lead to increased toxicity. These data highlight the importance of considering disease-specific biology when designing cytokine-targeting approaches, as strategies may not be broadly applicable across lymphoid and myeloid malignancies.

Our study focused on IFNγ because it is core to the initial burst of pro-inflammatory signaling by CAR T cells, and due to its pleiotropic effects on cells within the tumor microenvironment.

Further work is needed to more broadly define cytokine and cellular interactions within the CAR T/AML microenvironment and their impact on therapeutic response. Future studies in this area will guide the development of AML-specific CAR T cell strategies that enhance anti-leukemic activity without compromising CAR T cell function.

## Acknowledgements

The authors thank the University of Wisconsin Department of Pathology and Laboratory Medicine along with the Cellular and Molecular Pathology training program and NIH grant T32 GM135119 for funding and use of its facilities and services. The authors thank the University of Wisconsin Carbone Cancer Center (UWCCC) Flow Cytometry Laboratory, supported by P30 CA014520, for use of its facilities and services, including cell sorting performed on the BD FACSAria II Cell Sorter, supported by NIH Shared Instrumentation Grant 1S10RR025483-01. The authors also thank Dr. Crystal Mackall for generously providing many of the cell lines and CAR constructs used in this study. Illustrations in Figures 1,2,3 and S7, were created with Biorender.com.

## Authorship Contributions

NM designed and performed experiments, acquired and analyzed data, prepared the manuscript, and generated figures. IK, LM, JR, and OAK contributed to experimental work, data collection, and analysis. MC, SG, and LK assisted with experimental work and data acquisition. RMR conceptualized the study, supervised experimental design and data interpretation, acquired funding, and finalized the manuscript. All authors reviewed the manuscript and approved the final version.

## Reference List

1. Mitra A, Barua A, Huang L, Ganguly S, Feng Q, He B. From bench to bedside: the history and progress of CAR T cell therapy. Frontiers in Immunology. 2023;14.

2. Shahzad M, Nguyen A, Hussain A, et al. Outcomes with chimeric antigen receptor t-cell therapy in relapsed or refractory acute myeloid leukemia: a systematic review and meta-analysis. Frontiers in Immunology. 2023;14.

3. Cummins KD, Gill S. Chimeric antigen receptor T-cell therapy for acute myeloid leukemia: how close to reality? Haematologica. 2019/07/01;104(7).

4. Brudno JN, Kochenderfer JN. Toxicities of chimeric antigen receptor T cells: recognition and management. Blood. 2016;127(26):3321.

5. JC F, SL W, SL M, et al. Cytokine Release Syndrome After Chimeric Antigen Receptor T Cell Therapy for Acute Lymphoblastic Leukemia -PubMed. Critical care medicine. 2017 Feb;45(2).

6. Freyer CW, Porter DL. Cytokine release syndrome and neurotoxicity following CAR T-cell therapy for hematologic malignancies. Journal of Allergy and Clinical Immunology. 2020;146(5):940.

7. RQ L, L L, W Y, et al. FDA Approval Summary: Tocilizumab for Treatment of Chimeric Antigen Receptor T Cell-Induced Severe or Life-Threatening Cytokine Release Syndrome - PubMed. The oncologist. 2018 Aug;23(8).

8. Davila ML, Riviere I, Wang X, et al. Efficacy and Toxicity Management of 19-28z CAR T Cell Therapy in B Cell Acute Lymphoblastic Leukemia. Science translational medicine. 2014;6(224).

9. Gazeau N, Liang EC, Wu QV, et al. Anakinra for Refractory Cytokine Release Syndrome or Immune Effector Cell-Associated Neurotoxicity Syndrome after Chimeric Antigen Receptor T Cell Therapy. Transplantation and Cellular Therapy, Official Publication of the American Society for Transplantation and Cellular Therapy. 2023;29(7):430.

10. Norelli M, Camisa B, Barbiera G, et al. Monocyte-derived IL-1 and IL-6 are differentially required for cytokine-release syndrome and neurotoxicity due to CAR T cells. Nature Medicine. 2018;24(6):739.

11. Silveira CRF, Corveloni AC, Caruso SR, et al. Frontiers | Cytokines as an important player in the context of CAR-T cell therapy for cancer: Their role in tumor immunomodulation, manufacture, and clinical implications. Frontiers in Immunology. 2022/09/12;13.

12. Luciano M, Krenn PW, Horejs-Hoeck J. The cytokine network in acute myeloid leukemia. Frontiers in Immunology. 2022 Sep 28;13.

13. Epperly R, Gottschalk S, Velasquez MP. A Bump in the Road: How the Hostile AML Microenvironment Affects CAR T Cell Therapy. Frontiers in Oncology. 2020 Feb 28;10.

14. Bhagwat AS, Torres L, Shestova O, et al. Cytokine-mediated CAR T therapy resistance in AML. Nature Medicine. 2024;30(12):3697.

15. Wei Z, Xu J, Zhao C, et al. Prediction of severe CRS and determination of biomarkers in B cell-acute lymphoblastic leukemia treated with CAR-T cells. Frontiers in Immunology. 2023 Oct 3;14.

16. DT T, SF L, PA S, et al. Identification of Predictive Biomarkers for Cytokine Release Syndrome after Chimeric Antigen Receptor T-cell Therapy for Acute Lymphoblastic Leukemia - PubMed. Cancer discovery. 2016 Jun;6(6).

17. Bailey SR, Vatsa S, Larson RC, et al. Blockade or Deletion of IFNgamma Reduces Macrophage Activation without Compromising CAR T-cell Function in Hematologic Malignancies. Blood Cancer Discov. 2022;3(2):136–153.

18. Manni S, Del Bufalo F, Merli P, et al. Neutralizing IFNγ improves safety without compromising efficacy of CAR-T cell therapy in B-cell malignancies. Nature Communications. 2023;14(1):3423.

19. Johnson WT, Brumwell N, Ganesan N, et al. Emapalumab for the Treatment of Toxicities Associated with CD19 CAR-T Cells in Adult Patients. Transplantation and Cellular Therapy, Official Publication of the American Society for Transplantation and Cellular Therapy. 2025;31(2):S239.

20. Schuelke MR, Bassiri H, Behrens EM, et al. Emapalumab for the treatment of refractory cytokine release syndrome in pediatric patients. Blood Advances. 2023;7(18):5603.

21. Larson RC, Kann MC, Bailey SR, et al. CAR T cell killing requires the IFNγR pathway in solid but not liquid tumours. Nature. 2022;604(7906):563.

22. E T, S R, I K, et al. SOCS1 Protects Acute Myeloid Leukemia against Allogeneic T Cell-Mediated Cytotoxicity - PubMed. Blood cancer discovery. 05/05/2025;6(3).

23. Sayitoglu EC, Luca BA, Boss AP, et al. AML/T cell interactomics uncover correlates of patient outcomes and the key role of ICAM1 in T cell killing of AML. Leukemia. 2024;38(6):1246.

24. A M, M L, RT H, T T, T S, MM S. Antigen-dependent release of IFN-gamma by cytotoxic T cells up-regulates Fas on target cells and facilitates exocytosis-independent specific target cell lysis - PubMed. Journal of immunology (Baltimore, Md. : 1950). 07/01/2002;169(1).

25. Upadhyay R, Boiarsky JA, Pantsulaia G, et al. A critical role for fas-mediated off-target tumor killing in T cell immunotherapy. Cancer discovery. 2020 Dec 17;11(3).

26. Clement K, Rees H, Canver MC, et al. CRISPResso2 provides accurate and rapid genome editing sequence analysis. Nature Biotechnology 2019 37:3. 2019–02-26;37(3).

27. Sun Y, Wang S, Zhao L, Zhang B, Chen H. IFN-γ and TNF-α aggravate endothelial damage caused by CD123-targeted CAR T cell. OncoTargets and therapy. 2019 Jun 24;12.

28. RM R, F Z, KA F, et al. NOT-Gated CD93 CAR T Cells Effectively Target AML with Minimized Endothelial Cross-Reactivity - PubMed. Blood cancer discovery. 09/16/2021;2(6).

29. M M, RH R, H T, MK B. A T-cell-directed chimeric antigen receptor for the selective treatment of T-cell malignancies - PubMed. Blood. 08/20/2015;126(8).

30. D K, AC F, T N, et al. Intra-Leukemic Interferon Signaling Suppresses Expansion and Mediates Chemoresistance in Human AML - PubMed. Blood cancer discovery. 01/12/2026;7(1).

31. Wang B, Reville PK, Yassouf MY, et al. Comprehensive characterization of IFNγ signaling in acute myeloid leukemia reveals prognostic and therapeutic strategies. Nature Communications. 2024;15(1).

32. Vadakekolathu J, Minden MD, Hood T, et al. Immune landscapes predict chemotherapy resistance and immunotherapy response in acute myeloid leukemia. Science Translational Medicine. 2020-06-03;12(546).

33. Rimando J, Rettig MP, Christopher M, et al. Flotetuzumab and Other Cellular Immunotherapies Upregulate MHC Class II Expression on Acute Myeloid Leukemia Cells in Vitro and In Vivo. Blood. 2020/11/05;136(Supplement 1).

34. Linder A, Nixdorf D, Kuhl N, et al. STING activation improves T-cell-engaging immunotherapy for acute myeloid leukemia. Blood. 2025/05/08;145(19).

35. RG M, SP R, E S, et al. Tuning the Antigen Density Requirement for CAR T-cell Activity - PubMed. Cancer discovery. 2020 May;10(5).

36. Arcangeli S, Rotiroti MC, Bardelli M, et al. Balance of Anti-CD123 Chimeric Antigen Receptor Binding Affinity and Density for the Targeting of Acute Myeloid Leukemia. Molecular Therapy. 2017 May 4;25(8).

37. Kantari-Mimoun C, Barrin S, Vimeux L, et al. CAR T-cell Entry into Tumor Islets Is a Two-Step Process Dependent on IFNγ and ICAM-1. Cancer Immunology Research. 2021/12/01;9(12).

38. L M-L, A A, J P. How Do Cytotoxic Lymphocytes Kill Cancer Cells? - PubMed. Clinical cancer research : an official journal of the American Association for Cancer Research. 11/15/2015;21(22).

39. Korell F, Berger TR, Maus MV. Understanding CAR T cell-tumor interactions: Paving the way for successful clinical outcomes. Med. 2022/08/12;3(8).

40. Hong LK, Chen Y, Smith CC, et al. CD30-Redirected Chimeric Antigen Receptor T Cells Target CD30+ and CD30− Embryonal Carcinoma via Antigen-Dependent and Fas/FasL Interactions. Cancer Immunology Research. 2018/10/01;6(10).

41. Reina M, Espel E. Role of LFA-1 and ICAM-1 in Cancer. Cancers. 2017 Nov 3;9(11).

42. Bell M, Gottschalk S. Frontiers | Engineered Cytokine Signaling to Improve CAR T Cell Effector Function. Frontiers in Immunology. 2021/06/04;12.

43. Yeku OO, Purdon TJ, Koneru M, et al. Armored CAR T cells enhance antitumor efficacy and overcome the tumor microenvironment. Scientific Reports 2017 7:1. 2017–09-05;7(1).

