## Supplemental Figures and Table for "Disruption of the interferon-gamma axis limits chimeric antigen receptor T cell efficacy against acute myeloid leukemia"

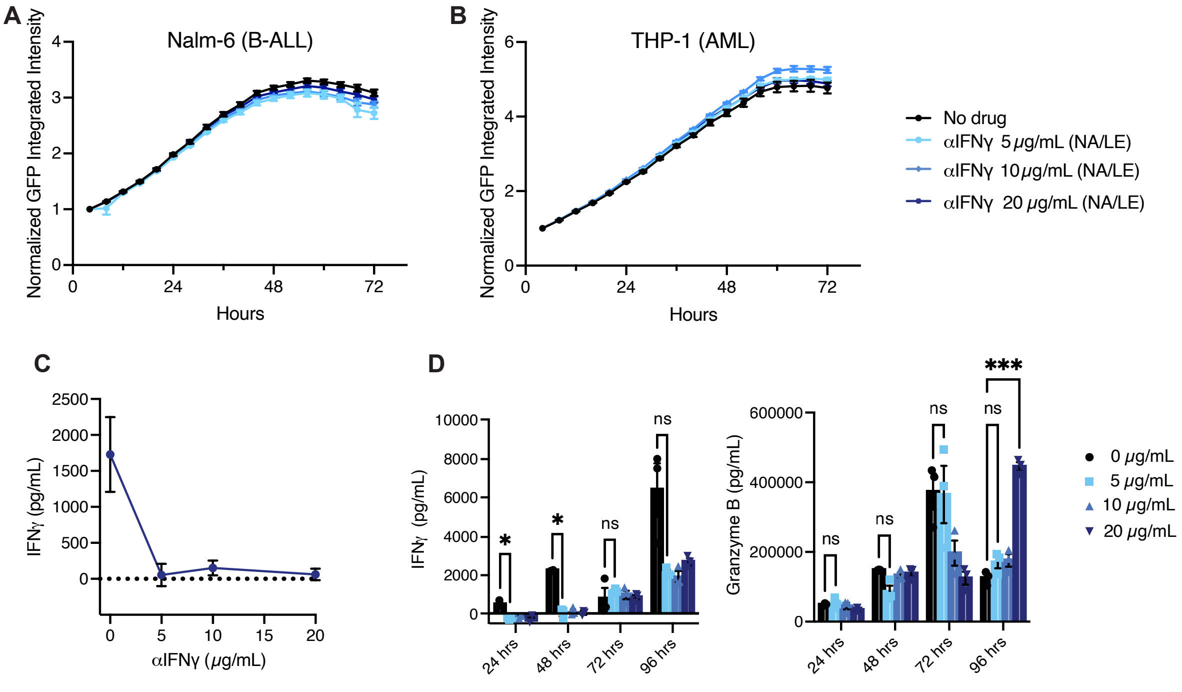


***Supplemental Figure 1. IFNγ blockade with αIFNγ is effective and non-toxic. A-B****. Nalm-6 (A) and THP-1 (B) cells, untreated or treated with azide-free/low endotoxin (NA/LE) αIFNγ at 5, 10, or 20 μg/mL. Tumor cell growth was monitored over 72 h by relative GFP intensity using Incucyte® live cell imaging.* ***C.*** *ELISA quantification of IFNγ concentration in CD123 CAR and THP-1 co-culture supernatants collected after 24 h with increasing doses of αIFNγ (NA/LE)* ***D.*** *ELISA quantification of IFNγ (left) and Granzyme B (right) in serial stimulation co-culture supernatants collected every 24 h for 96 h, n=3 T cell donors (two-way ANOVA).*

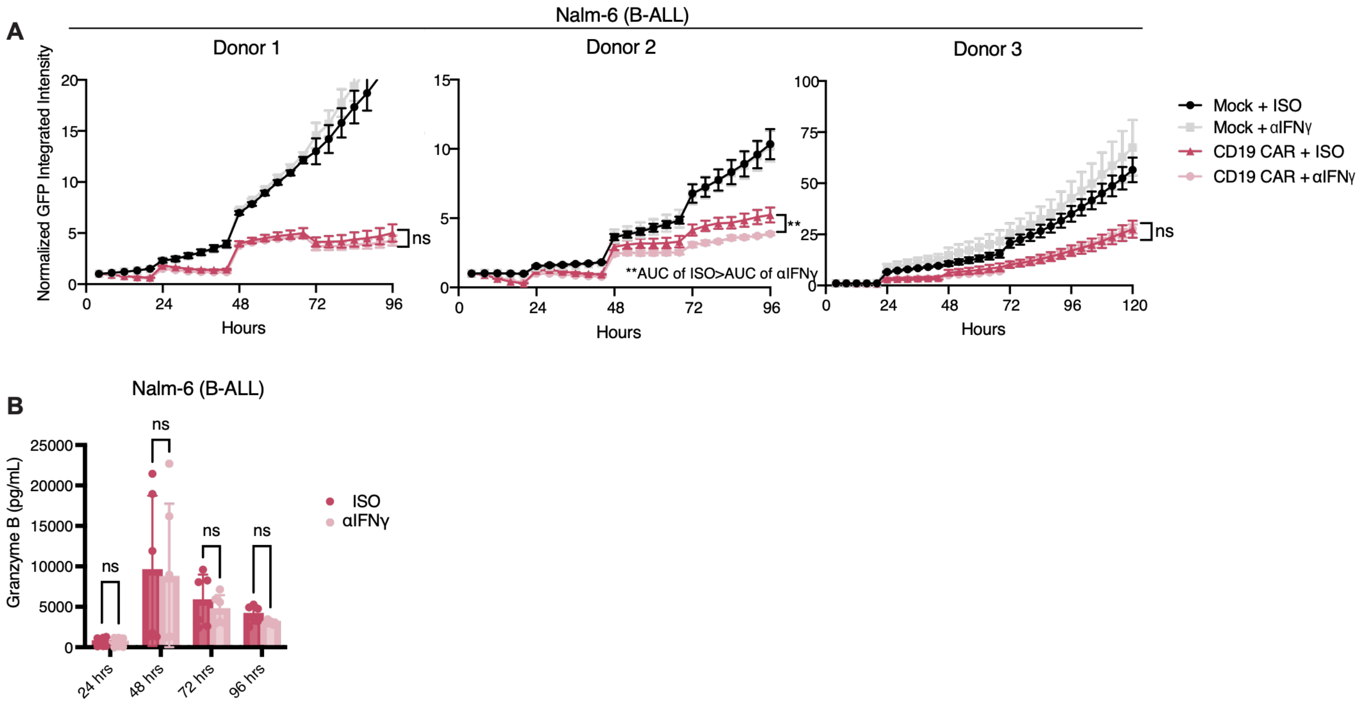


***Supplemental Figure 2. IFNγ blockade does not inhibit CD19 CAR T cell cytotoxicity of B-ALL. A.*** *Individual T cell donor replicates in serial stimulation co-culture of Nalm-6 cells and CD19 CAR T cells with or without $\alpha$IFN$\gamma$ at a 1:4 effector to target (E:T) ratio. Tumor growth was assessed as GFP integrated intensity normalized from hour 4. Statistical comparisons differences between CAR+ISO and CAR+$\alpha$IFNγ were performed for each individual donor, n=3 T cell donors (Welch's t-test of AUC).* ***B.*** *Granzyme B concentrations in Nalm-6/CD19 CAR co-culture supernatants measured by ELISA every 24 h over 96 h at 1:4 E:T, n=3 T cell donors (two-way ANOVA).*

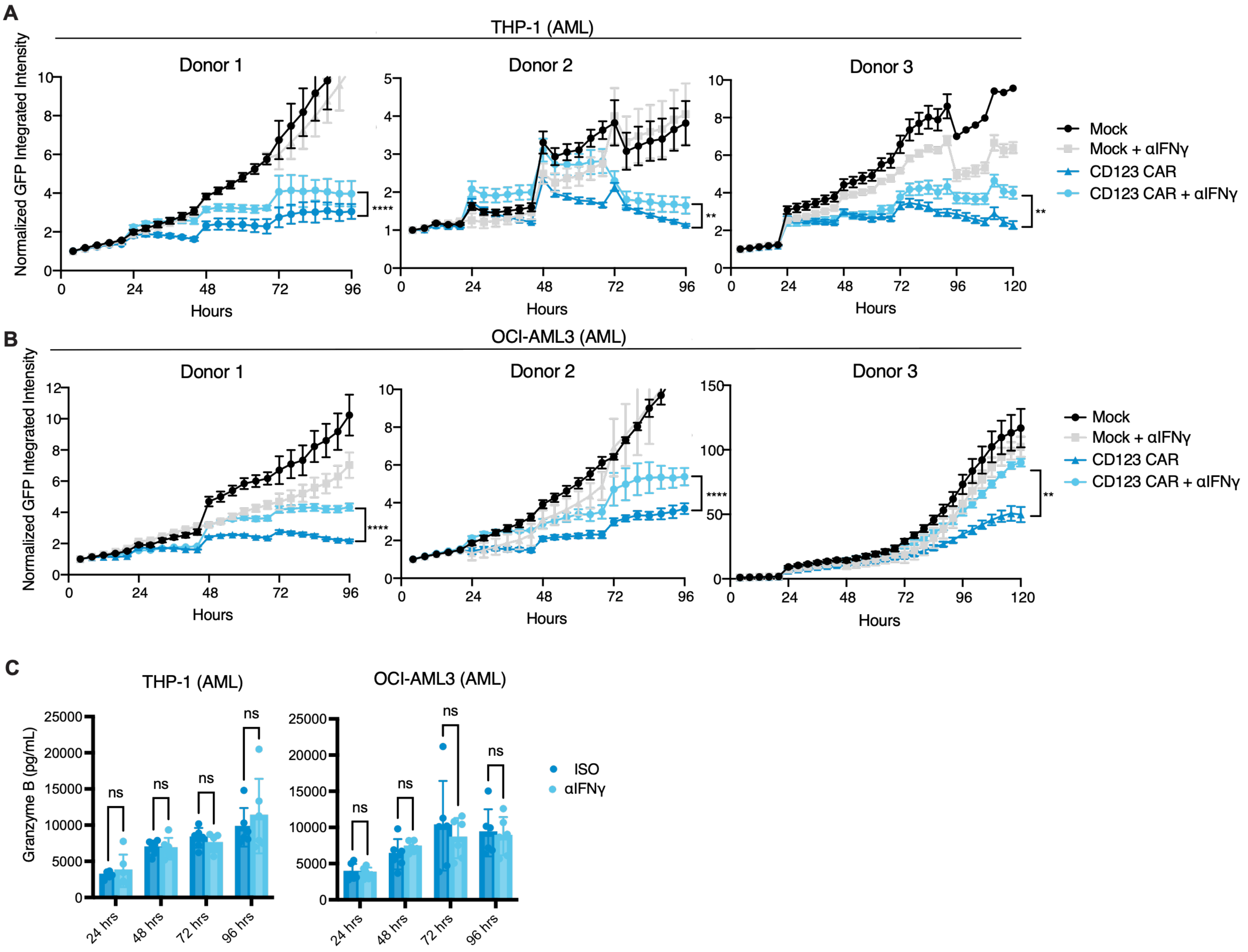


***Supplemental Figure 3. IFNγ blockade does inhibit CD123 CAR T cell killing of AML. A-B.*** *Individual T cell donor replicates in serial stimulation co-culture of (A) THP-1 or (B) OCI-AML3 cells and CD123 CAR T cells with or without αIFNγ at 1:4 E:T ratio. Tumor growth was assessed as GFP integrated intensity normalized to values at 4 h. Statistical comparisons on differences between CAR+ISO and CAR+αIFNγ were performed for each individual donor, n=3 T cell donors (Welch's t-test of AUC).* ***C.*** *Granzyme B concentrations in CD123 CAR and (left) THP-1 and (right) OCI-AML3 co-culture supernatants measured by ELISA every 24 h over 96 h at 1:4 E:T, n=3 T cell donors (two-way ANOVA).*

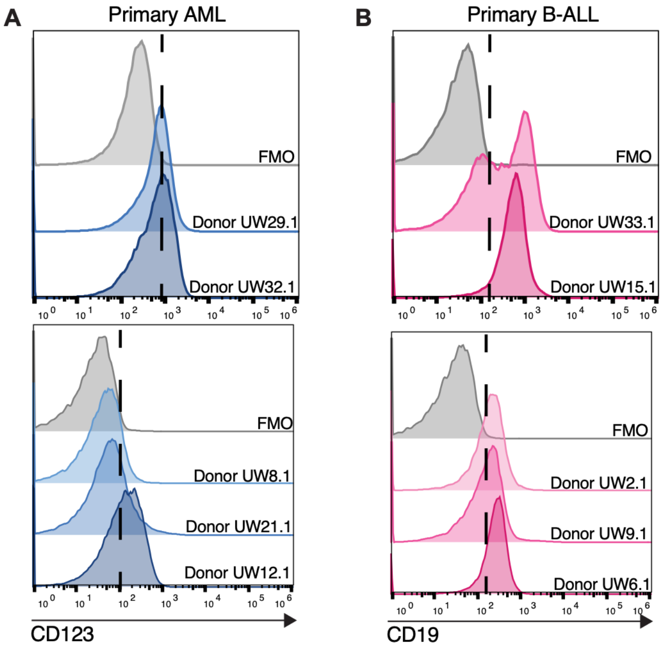


***Supplemental Figure 4. Target antigen density on primary patient-derived leukemia. A-B.*** *Flow cytometry plots of (A) CD123 or (B) CD19 expression on the surface of each primary (A) AML or (B) B-ALL patient sample. Dotted line was drawn based off the fluorescence minus one (FMO) control for each respective flow cytometry run.*

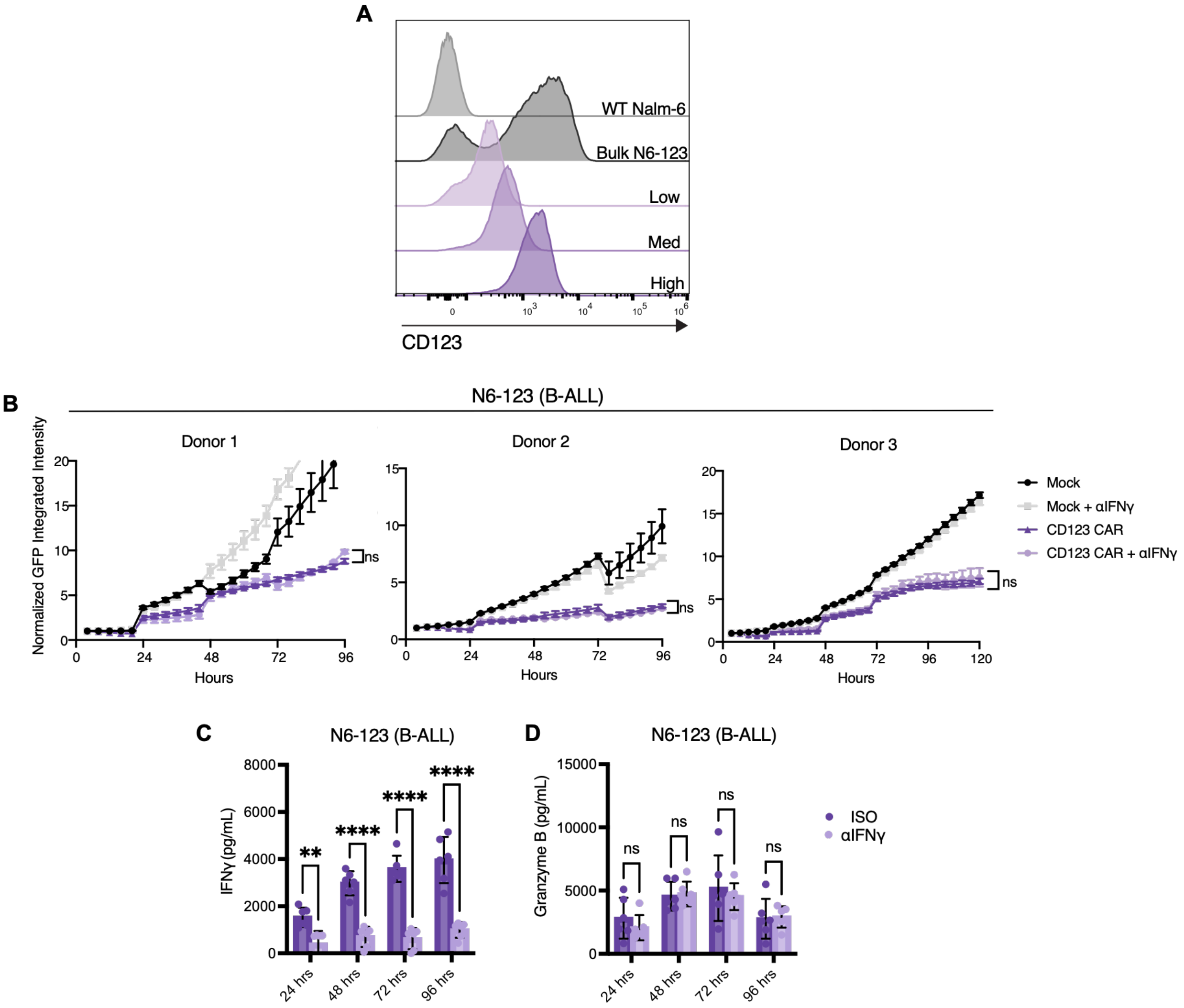


***Supplemental Figure 5. IFNγ blockade does not impact CD123 CAR T cell killing of an engineered B-ALL cell line.*** ***A.*** *Flow cytometry of CD123 expression on Nalm6 cells engineered to express CD123 (N6-123), comparing WT N6 cells to the bulk engineered population to single cell sorted N6-123 lines.* ***B.*** *Individual T cell donor replicates in serial stimulation co-culture of N6-123 cells and CD123 CAR T cells with or without αIFNγ at a 1:4 E:T ratio. Tumor growth and statistical comparisons assessed as in Fig S2 and S3, n=3 T cell donors (Welch's t-test of AUC).* ***C-D.*** *ELISA quantification of IFNγ (C) and Granzyme B (D) in supernatant from N6-123 and CD123 CAR cocultures every 24 h for 96 h, n=3 T cell donors (two-way ANOVA).*

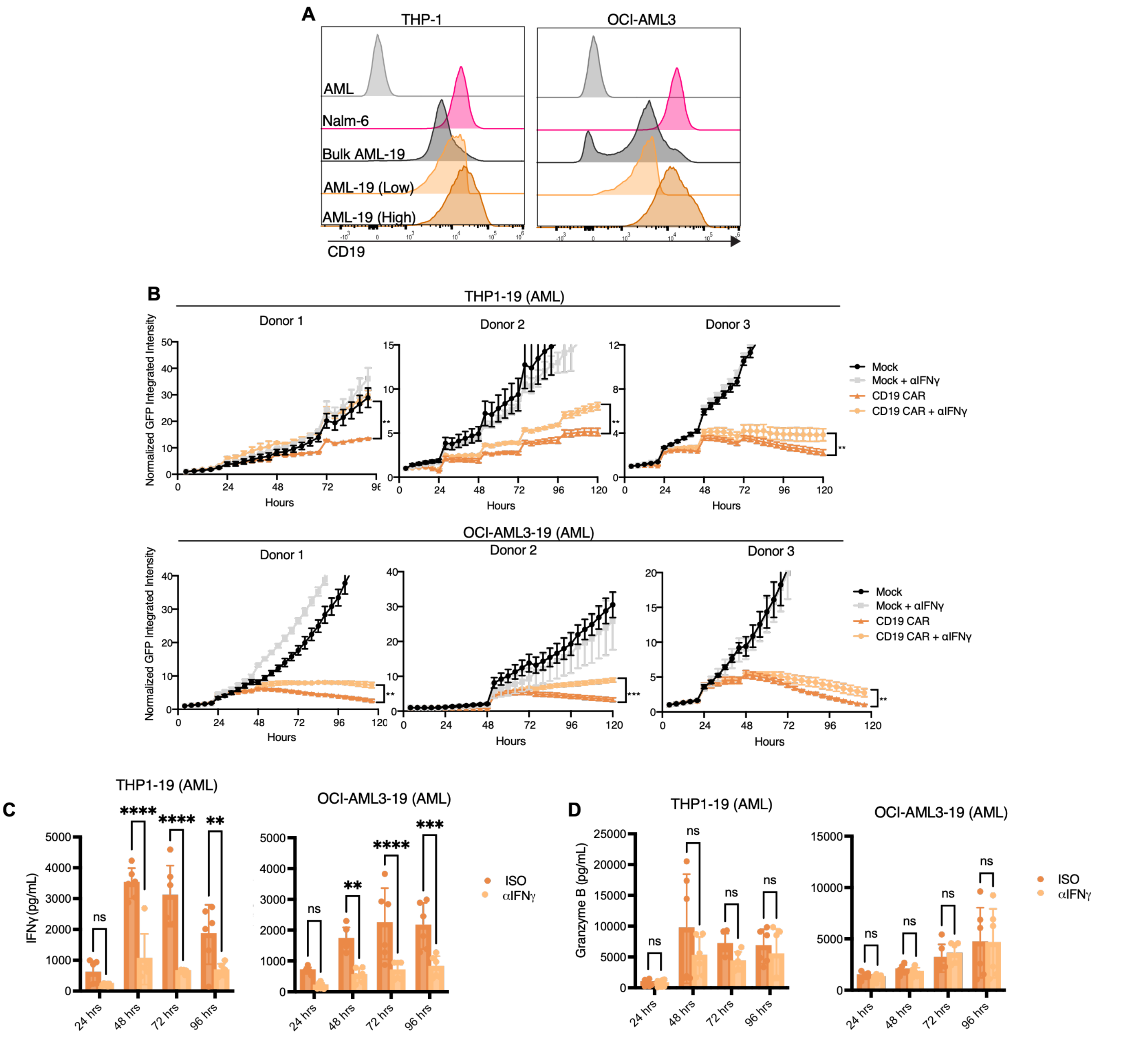


***Supplemental Figure 6. IFNγ blockade does inhibit CAR T cell killing of CD19-expressing AML. A.*** *Flow cytometry of CD19 expression on Bulk AML-19 cells and single cell sorted AML-19 lines as compared to WT Nalm-6 cells and WT AML.* ***B.*** *Individual T cell donor replicates in serial stimulation co-culture of (top) THP1-19 or (bottom) OCI-AML3-19 cells and CD19 CAR T cells with or without αIFNγ at 1:4 E:T. Tumor growth and statistical comparisons assessed as in previous figures, n=3 T cell donors. (Welch's t-test of AUC).* ***C-D.*** *ELISA quantification of IFNγ (C) and Granzyme B (D) in supernatant from AML-19 and CD19 CAR cocultures every 24 hours for 96 hours, n=3 (two-way ANOVA).*

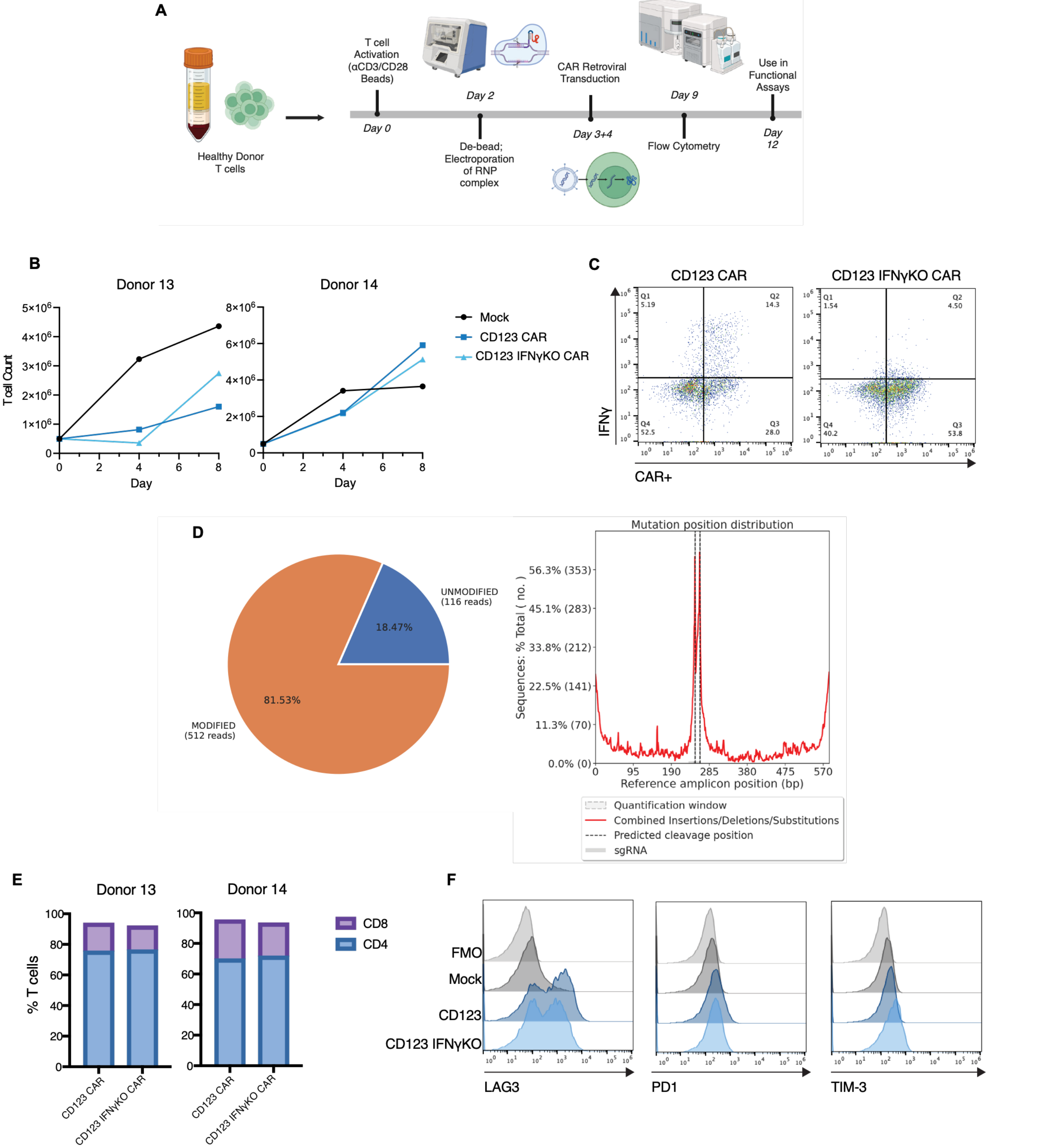


***Supplemental Figure 7. Validation and characterization of IFNγKO CAR T cells. A.*** *Schematic of the IFNγKO CAR T cell manufacturing process.* ***B.*** *CAR T cell expansion plots with live cell counts every 4 days post-electroporation.* ***C.*** *Intacellular flow cytometry of IFNγ (y-axis) versus CAR positivity (x axis) assessed 4 h post activation with PMA + ionomycin.* ***D.*** *Genome editing sequence analysis of IFNγKO CAR T cells with quantification of edited reads (left) and mutation distribution relative to sgRNA cutsites (right). DNA amplicons of the cut region were generated with PCR and sequenced by Plasmidsaurus. Genome editing analysis was run with CRISPResso2.* ***E.*** *Percentage CD4 and CD8 positive T cells on WT and IFNγKO CD123 CAR T cells as measured by flow cytometry.* ***F.*** *Surface expression of LAG-3, PD1, and TIM-3 on WT and IFNγKO CD123 CAR T cells as compared to mock T cells and FMO controls.*

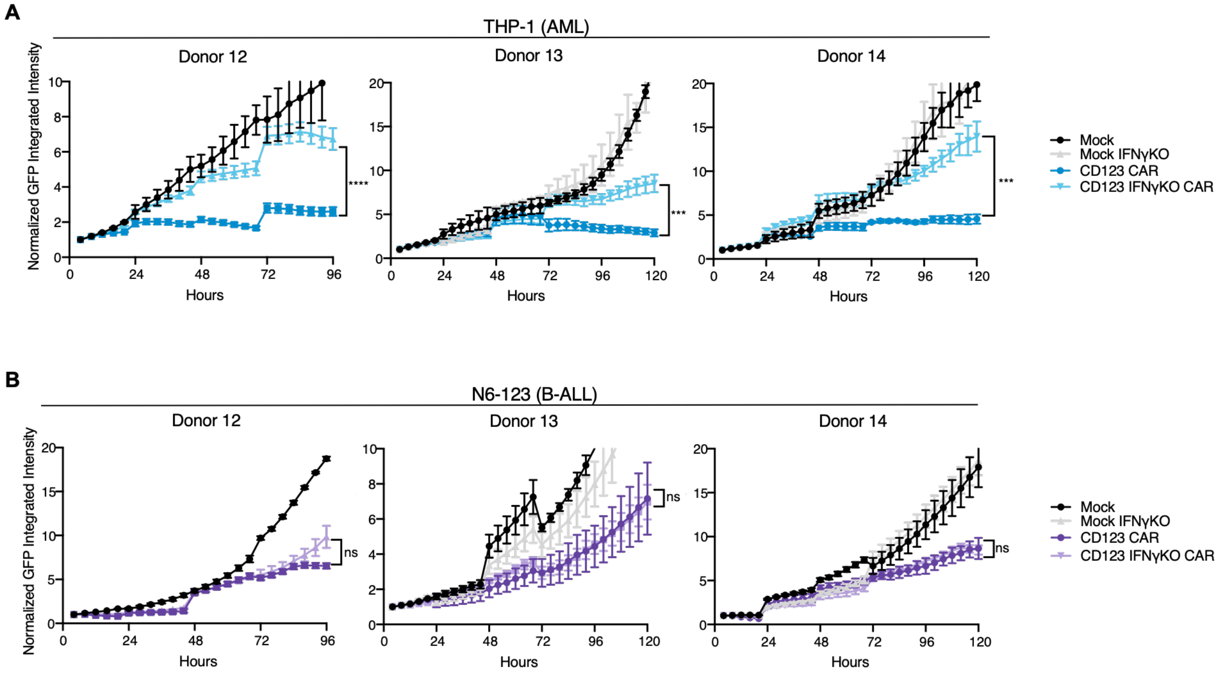


***Supplemental Figure 8. IFNγKO inhibits CD123 CAR T cell killing of AML but not B-ALL. A-B.*** *Individual T cell donor replicates in serial stimulation co-culture of (A) THP-1 or (B) N6-123 cells and CD123 CAR T cells with or without IFNγKO at a 1:2 E:T. Tumor growth and statistical comparisons assessed as in previous figures, n=3 T cell donors (Welch's t-test of AUC).*

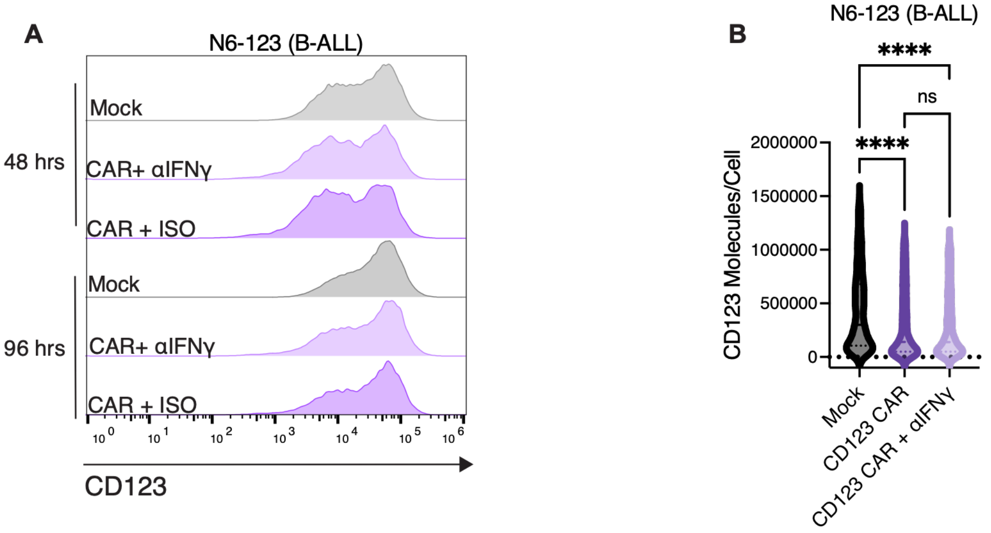


| | *CD123 Molecules/Cell* | |
|---|---|---|
| *Condition* | *Mean (SD)* | *Median (IQR)* |
| Mock | 543146 (575822) | 360032 (117434-775968) |
| CAR + ISO | 394861 (500998) | 167768 (56297-607313) |
| CAR + αIFNγ | 399259 (519554) | 168016 (56167-586138) |

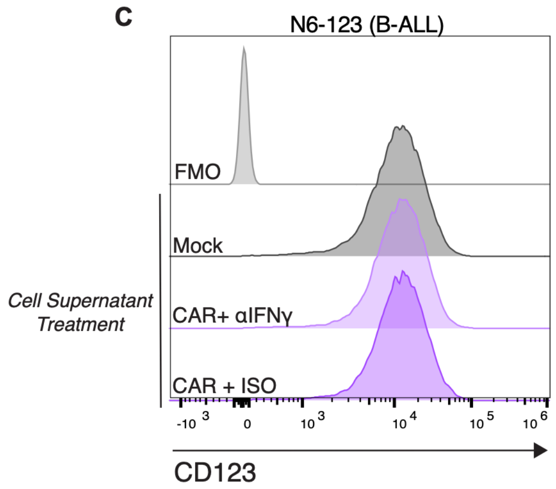


***Supplemental Figure 9. CD123 on N6-123 cells is not upregulated the context of CAR T cell inflammation. A-B.*** *Flow cytometry of CD123 expression on N6-123 cells at 48 and 72 h during a serial stimulation assay, with or without αIFNγ treatment at 1:16 E:T represented as (A) fluorescent intensity histograms, (B) molecules per cell calculated same as in Fig 4B also on 1x10$^4$ individual cells (two-way ANOVA).* ***C.*** *Flow cytometry of CD123 expression on N6-123 cells exposed to supernatant from N6-123/Mock, N6-123/CAR, or N6-123/CAR+αIFNγ co-cultures for 24 h.*

.

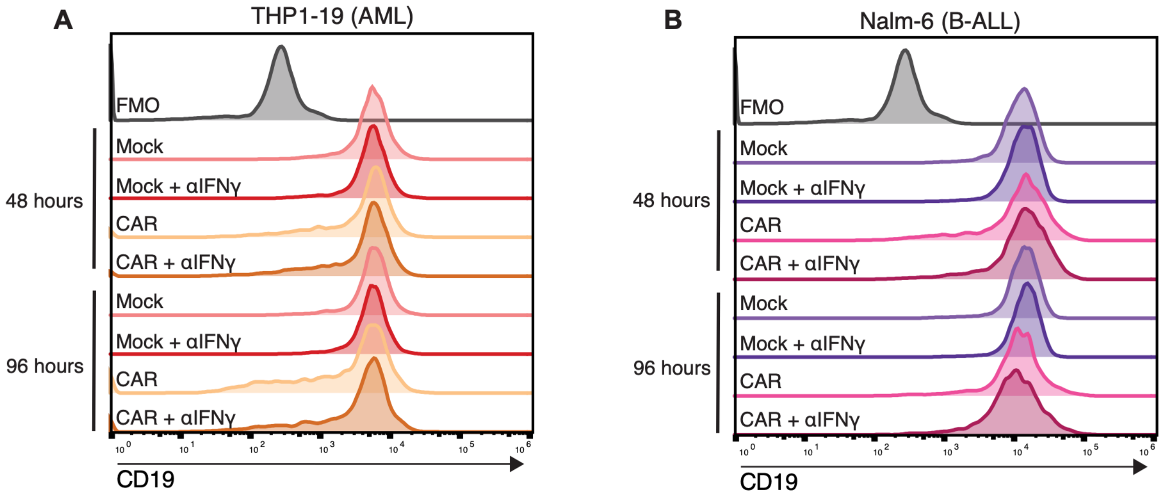


***Supplemental Figure 10. CD19 expression on B-ALL and engineered AML with IFNγ blockade. A.*** *Flow cytometry of CD19 expression on (A) THP1-19 and (B) Nalm-6 cells at 48 and 72 h during a serial stimulation assay, with or without αIFNγ treatment represented as fluorescent intensity histograms.*

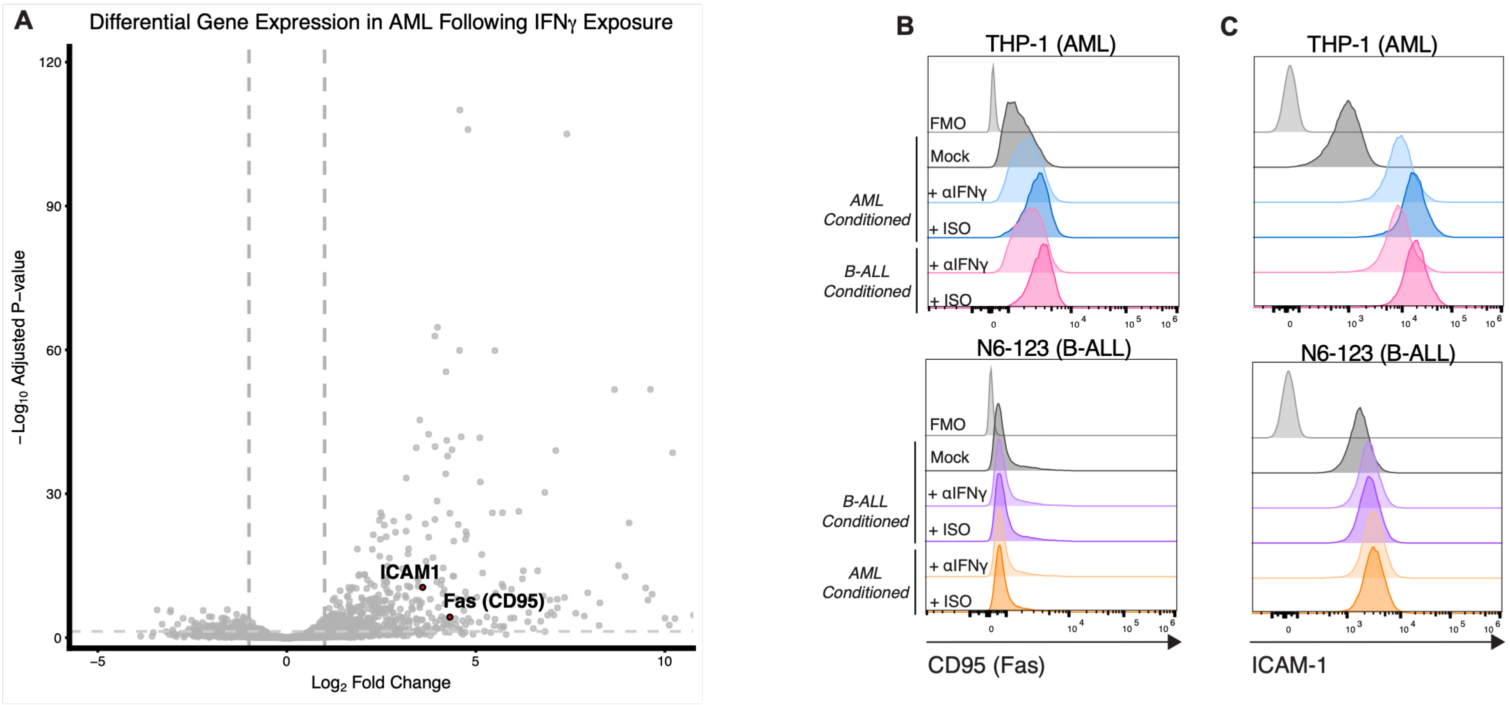


***Supplemental Figure 11. ICAM-1 and Fas expression on tumor cell lines. A.*** *Volcano plot showing differentially expressed genes in AML cell lines (n=3) following 24-hour exposure to IFNγ, analyzed using a DESeq2 model accounting for both cell line and treatment condition (design = ~ cell_line + condition). Each point represents an individual gene plotted by log2 fold change (x-axis) and −log10(p-value) (y-axis). Genes significantly upregulated or downregulated following IFNγ exposure are distributed across the plot, with ICAM1 and FAS highlighted in red as key IFNγ-responsive genes implicated in immune synapse formation and apoptosis signaling, respectively. Horizontal dashed line indicates p = 0.05 significance threshold; vertical dashed lines indicate log2 fold change thresholds of ±2. (GEO accession GSE159991).* ***B-C.*** *Flow cytometry of (B) Fas and (C) ICAM-1 expression on (top) THP-1 or (bottom) N6-123 cells after 24 h of exposure to conditioned media from THP-1 or N6-123 co-cultures (+ mock T cells, CD123 CAR T cells + isotype control, or CD123 CAR T cells + αIFNγ).*

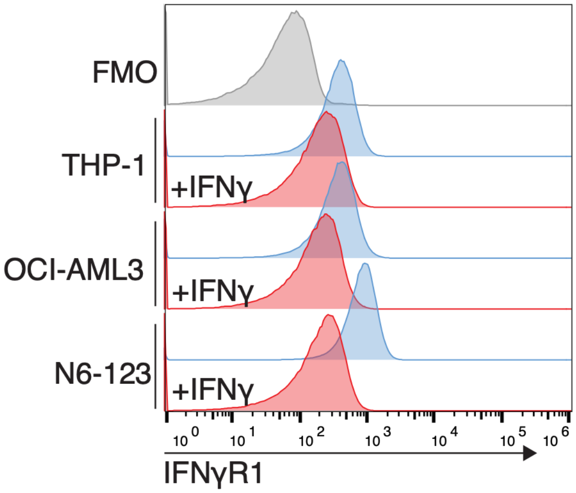


***Supplemental Figure 12. Exogenous IFNγ added to AML and B-ALL cell lines.*** *Flow cytometry analysis of IFNγR1 expression on AML and B-ALL cell lines after 24 h pre-incubation with 10 ng/mL of IFNγ.*

| Antibody | Fluorophore | Manufacturer | Catalog # |
|---|---|---|---|
| Anti-human IFNγ | PE | Biolegend | 506506 |
| Anti-human CD8 | PE-Cy7 | BD Biosciences | 557746 |
| Anti-human CD4 | BV711 | BioLegend | 317440 |
| Anti-human PD-1 | BV605 | BioLegend | 329924 |
| Anti-human TIM-3 | BV421 | BioLegend | 345013 |
| Anti-human ICAM-1 | PE | BioLegend | 322706 |
| Anti-human CD95 (Fas) | BV711 | BioLegend | 305640 |
| Anti-human CD123 | PE-Cy7 | BioLegend | 306026 |
| Anti-human CD19 | APC-Vio770 | Miltenyi Biotec | 130-113-643 |
| Anti-human LAG-3 | PE | BioLegend | 369306 |
| Goat anti-mouse IgG $F(ab')_2$ | AF647 | Jackson ImmunoResearch | 115-606-072 |

***Supplementary Table 1. List of fluorophores and antibodies utilized in flow cytometry***
