## Supplemental Methods for "Disruption of the interferon-gamma axis limits chimeric antigen receptor T cell efficacy against acute myeloid leukemia"

*Cell Lines*

Packaging cell line 293T stably transduced with retroviral gag and pol proteins (293GP), as well as tumor cell lines THP-1, OCI-AML3, and Nalm-6 cells stably transduced with GFP-luciferase were kindly provided by Dr. Crystal Mackall (Stanford University). N6-123, THP1-19, and OCI-AML3-19 cell lines were generated in the Richards Lab with retroviral transduction as described below. All cell lines were verified by short tandem repeat analysis and confirmed to be *Mycoplasma* negative by PCR at least every 6 months.

*CAR Constructs*

The CAR construct containing an scFv targeting CD123 (26292, patent# WO2017075147A1)[27] cloned as previously described[28] into a MSGV1 backbone containing the CD28-CD28-CD3ζ hinge, transmembrane, and intracellular signaling domains. The CD19-28ζ CAR was subcloned by a restriction enzyme digest the of CD19 scFv sequence (FMC63, patent #US9701758B2) from a construct containing the CD8α-4-1BB-CD3ζ hinge, transmembrane, and intracellular signaling domains, followed by a ligation into the CD28-CD28-CD3ζ MSGV1 backbone. Constructs with CD28-CD28-CD3ζ or CD8α-4-1BB-CD3ζ backbones were provided by Dr. Crystal Mackall.

*Primary T cell Isolation*

Human peripheral blood mononuclear cells (PBMCs) were isolated from whole blood or leukocyte reduction system cones (Versiti) from healthy donors. T cells were separated with Lymphoprep (STEMCELL Technologies, Cambridge, MA, USA; Cat#18061) and isolated using either the EasySep Human T cell Isolation Kit (STEMCELL Technologies; Cat#17951) or RosetteSep Human T cell Enrichment Cocktail (STEMCELL Technologies; Cat#15061) according to manufacturer instructions. Isolated T cells were frozen down in CryoStor CS10 (STEMCELL Technologies; Cat#100-1061) medium and stored at -150°C for later use. Processing of healthy donor PBMCs was done under UW-Madison IRB approval (I2017-1070).

*Retrovirus Production*

Retrovirus encoding the CD123 CAR, CD19 CAR, full length CD123, or truncated CD19 (tCD19, containing only the extracellular and transmembrane domains) were generated by transient transfection of 293GP cells with the plasmid encoding the CAR or surface protein and the RD114 envelope protein using Lipofectamine 2000 (Invitrogen, Carlsbad, CA, USA; Cat#11668027). Viral supernatants were collected at 48 and 72 h post-transfection, centrifuged at 2000 xg for 10 min, then either used fresh or stored at -80°C for future use.

*Bulk-RNA Sequencing Analysis*

RNASeq was performed on three AML cell lines (Kasumi-1, NOMO-1, THP-1), either at baseline or with exposure to pro-inflammatory cytokines including IFNγ, as described previously[28]. The dataset had been deposited in NCBI Gene Expression Omnibus (GEO) and was accessible with the accession number GSE159991. Differential gene expression analysis comparing unstimulated versus IFNγ exposed AML cells was performed using DESeq2 accounting for both cell line and treatment condition (design = ~ cell_line + condition). Differentially expressed genes were filtered in R using a statistical filter of false discovery rate (FDR) <0.05 and log2fold change > 2. Gene expression levels were visualized using the ggplot2 package in R.
